# A teneurin-3 microphthalmia mutation disrupts *trans* adhesion for specific alternative splicing isoforms

**DOI:** 10.1101/2025.02.28.640769

**Authors:** Christos Gogou, Stefan Loonen, Maja Napieraj, Cátia P Frias, Nikolina Šoštarić, Dimphna H Meijer

**Affiliations:** Department of Bionanoscience, Kavli Institute of Nanoscience Delft, Delft University of Technology, van der Maasweg 9, 2629HZ Delft, the Netherlands

**Keywords:** neuronal recognition, cell adhesion, alternative splicing, microphthalmia, structural biology

## Abstract

Microphthalmia (MCOP) is a developmental eye disorder in which one or both eyes are abnormally small. This condition is often accompanied by cognitive impairments. Teneurin-3, a cell adhesion molecule with important functions in axon pathfinding and synaptic organization, has been repeatedly implicated with MCOP, suggesting its potential role in the etiology of this complex developmental disorder. It was previously shown that two small alternatively spliced exons – inserts A and B – instigate large structural reorganizations of the teneurin-3 covalent homodimer. Here, we map the MCOP mutation R2579W to an intramolecular interface specific to splice insert A-containing isoforms. We demonstrate that the MCOP mutation leads to a loss of transcellular adhesion, predominantly affecting these isoforms. Using small-angle X-ray scattering we establish that, despite these functional disruptions, the structural compactness of all mutant isoforms is preserved. Molecular dynamics simulations predict a global stabilization of the A_1_ mutant molecule compared to its wildtype counterpart, due to changes in local compact dimer interactions around the mutated residue. We experimentally validate these findings with biophysical assays. With cryo-EM analysis, we confirm the overall compactness with reduced conformational flexibility and reveal that the mutant tryptophan binds to a hydrophobic pocket in the structure. With these data, we provide a model for how increased stability in *cis* disrupts *trans* oligomerization needed for cellular adhesion. Altogether, we demonstrate that the MCOP-associated mutation in teneurin-3 causally disrupts cellular adhesion in an isoform-specific manner, thereby likely directly contributing to the development of the disease in MCOP patients.

## Introduction

Microphthalmia (MCOP) is a congenital neurological disease that is typified by severely malformed eyes, often in combination with delayed cognitive development^1,2^. An estimated 14 per 100,000 newborns suffer from MCOP as either a stand-alone disease or as part of one of ∼200 syndromes^2–4^. Roughly 30% of the syndromic patients exhibit brain malformation, and a similar percentage displays intellectual disabilities^3^. According to the ClinVar database, 22 pathological missense mutations in various genes are associated with MCOP^5,6^. One such pathological missense mutation is the R2579W substitution in the *TENM3* gene on chromosome 4, which encodes teneurin-3 (Ten3) in humans. Aside from the Ten3 R2579W missense mutation listed in ClinVar^6^, various homozygous nonsense mutations in Ten3 emerged in multiple MCOP case reports^7–11^. In general, such nonsense mutations lead to truncated proteins that may be non-functional, unstable and are often quickly degraded in the cell^10^. Clinical mutations that do not result in truncation, degradation or mislocalization of the protein permit detailed molecular investigation of the disrupted function of gene products.

Cell adhesion molecules (CAM) on the neuronal plasma membrane act as guidance cues for axonal pathfinding during network formation in the brain^12–16^. Of the four teneurin family members of the type II transmembrane CAM family, Ten3 is involved in neuronal wiring of the hippocampus and visual system of different animal species^17–28^. In zebrafish, Antinucci and colleagues found that reducing Ten3 expression levels leads to improper formation of the inner plexiform layer of the retina and caused defects in connections between retinal ganglion cells (RGCs) and their target area, the tectal neuropil^19^. As a result, the fish had difficulty responding to certain directional and orientational inputs of visual stimuli. In Ten3 knockout mice, defects were observed in ipsilateral projections between RGCs and the dorsal lateral geniculate nucleus (dLGN), leading to difficulties in tasks that require binocular vision^25,26^. More recently, miswiring in Ten3 knockout mice was also observed between a light-sensitive subtype of RGCs (the intrinsically photosensitive RGCs or ipRGCs) and the suprachiasmic nucleus (SCN)^23^. At the molecular level, a combination of repulsive interactions with adhesion G protein-coupled receptor latrophilin (Lphn) and attractive homophilic interactions act as guidance cues for the proper wiring^22^. For instance, in the mouse hippocampus, Ten3-expressing axons from the proximal CA1 area project across the Lphn-expressing distal CA1 and proximal subiculum, over to the distal subiculum where Ten3 is expressed again^21,22^. *In vitro* studies with retinal neurons have recapitulated these attractive homophilic interactions, showing that Ten3-expressing neurons from chicken retinal explants preferentially grew on substrates coated with purified Ten3^29^. Altogether, across species, Ten3 is crucial for the proper connectivity of the visual system and the hippocampus in the brain. Pathogenic mutations such as the MCOP-associated R2579W substitution may thus directly interfere with Ten3’s wiring of these regions, as suggested by the clinical manifestation of visual impairment for this particular mutation.

Structural biology studies revealed that teneurins are composed of a large globular superfold at the extracellular side comprising multiple domains, starting from an N-terminal calcium-binding C-rich domain, a transthyretin-like (TTR) domain, a fibronectin domain (FN-plug), and followed by an NHL β-propeller^30–34^. The tyrosine- and aspartate-rich (YD) shell is the central and largest domain around which the other domains are stabilized. An additional linker domain threads the YD shell which upon exit is followed by the antibiotic-binding domain (ABD) and Tox-GHH-like domain. This full assembly is attached to a single-pass transmembrane helix via an immunoglobulin (Ig) fold and eight flexible repeats of an endothelial growth factor (EGF)-like domain. Importantly, teneurin-3 is a constitutive *cis* dimer in which the monomers are linked by two disulfide bridges between EGF repeats 2 and 5^35,36^. Many of the different domains are found to be involved in intramolecular contacts across teneurin homologues, as well as in intermolecular contacts in *cis* and in *trans* configurations with teneurins and other proteins^29–34,37^. The YD shell, for instance, harbours a heterophilic binding site for Lphn^32,33,38^.

The extracellular domain (ECD) of Ten3 contains two alternatively spliced regions: splice inserts A between EGF7-EGF8, and B within the NHL domain. The resulting splicing isoforms are referred to as A_0_B_0_ (both inserts absent), A_1_B_1_ (both present), and A_0_B_1_ and A_1_B_0_ (insert B or A present, respectively). All isoforms exist both in mice and humans^39^. Berns and colleagues revealed differential expression of these four isoforms in different parts of mouse hippocampus^21^. All isoforms were expressed in the subiculum, whereas A_0_B_0_ was not detected in the CA1 region. Spatial control of their expression hints at splicing-dependent roles within the brain, but such isoform-specific functions are not resolved. We previously solved the structures of the four different isoforms, which provided the mechanistic basis for the splicing-dependent differences in transcellular adhesion^29^. Shortly, in absence of splice insert A, splice insert B instigates a global reorganization of the superfolds within the ECD, where each isoform forms a different ‘compact dimer’ contact between the two superfolds. The presence of splice insert A prompts a third compact dimer configuration, which is not affected by the presence or absence of splice insert B. All compact dimer interfaces thus constitute additional non-covalent interactions between the two monomers in the EGF-linked constitutive dimer.

In this work, we map the MCOP-associated R2579W to an EGF8-ABD intramolecular compact dimer interface that is specific to the A_1_ isoforms. We use *in vitro* cell clustering assays to show that the MCOP mutation causes a loss of transcellular clustering for A_1_ isoforms, but not for A_0_ isoforms. Ten3 thereby appears to be a unique case in which a missense mutation present in all isoforms only affects a specific subset of isoforms. Despite these functional differences between the wildtype (WT) and pathological mutant proteins, the compact dimer conformations are conserved in all isoforms, as shown by small-angle X-ray scattering. Molecular dynamics (MD) simulations show that the tryptophan substitution locally interacts with a different set of residues than the wildtype arginine, thereby stabilising the compact dimer. Thermal stability- and size-exclusion chromatography-based assays confirm compact *cis* dimer stabilization. Finally, cryo-electron microscopy (cryo-EM) reconstruction of the stabilized A_1_B_1_ isoform confirms preservation of the compact dimer conformation and captures the hydrophobic contact that was predicted by the MD simulations. A 3D variability analysis of the cryo-EM particle set additionally reveals a restricted conformational freedom of the stabilized mutant compact dimer. Our data indicate that mutagenic stabilization of the *cis* compact dimer conformation in A_1_ isoforms inhibits *trans* interactions across cellular junctions for these Ten3 variants, specifically. These structural and functional insights shed light on the underlying mechanistic basis of microphthalmia and may, for instance, pave the way for improved diagnostics of this disease.

## Results

### R2579W mutation maps to A_1_-specific compact dimer interface of teneurin-3

Recent studies have identified various mutations in the teneurin-3 (Ten3) gene in patients suffering from microphthalmia. These mutations, except for R2579W (corresponding to A_1_B_1_ amino acid numbering; R2563W, R2570W, and R2572W in A_0_B_0_, A_0_B_1_, and A_1_B_0_ respectively), are nonsense mutations or induce frameshifts followed by premature stop codons, resulting in truncated proteins (Fig. 1a). Truncations at these specific sites result in a near-complete loss of the superfold (Fig. 1b and Supplementary Fig. 1a), the domain carrying multiple protein-protein interaction surfaces with functional importance^29,30,32–34^. The R2579W mutation, however, does not result in a truncated protein, but instead constitutes a single amino acid mutation that may affect the function of the full-length protein. This provides a unique opportunity to study the molecular mechanisms underlying both normal physiological processes and pathology. The patient in which the R2579W mutation was uncovered also possessed an additional Ten3 mutation, A1365G, classifying it as compound heterozygous (Fig. 1a)^11^. However, since this male patient’s brother and father also carried the A1365G mutation, but did not display any disease symptoms, we focus on R2579W. The Ten3 MCOP mutation localizes in one of the loops of the ABD domain, namely L2578-S2582 (Fig. 1c, on the right). Previous high-resolution structural analysis revealed that this loop and the β-strand directly flanking it (G2583-A2586) are unstructured in the compact dimer conformations of the splice variants that harbour splice insert A (A_1_B_0_ and A_1_B_1_, Fig. 1c)^29^. Displacement of this region renders the ABD accessible for the EGF8 binding required to form the A_1_ compact dimer. Negative stain structural analysis of the isoforms that lack insert A (A_0_B_0_ and A_0_B_1_) predicts that the MCOP-containing loop, and the β-strand directly flanking it, are located away from their respective intramolecular compact dimer interfaces^29^ (Fig. 1c). The organization of this loop may play an important role in the compact dimerization of the A_1_B_0_ and A_1_B_1_ isoforms, specifically.

**Figure 1:**
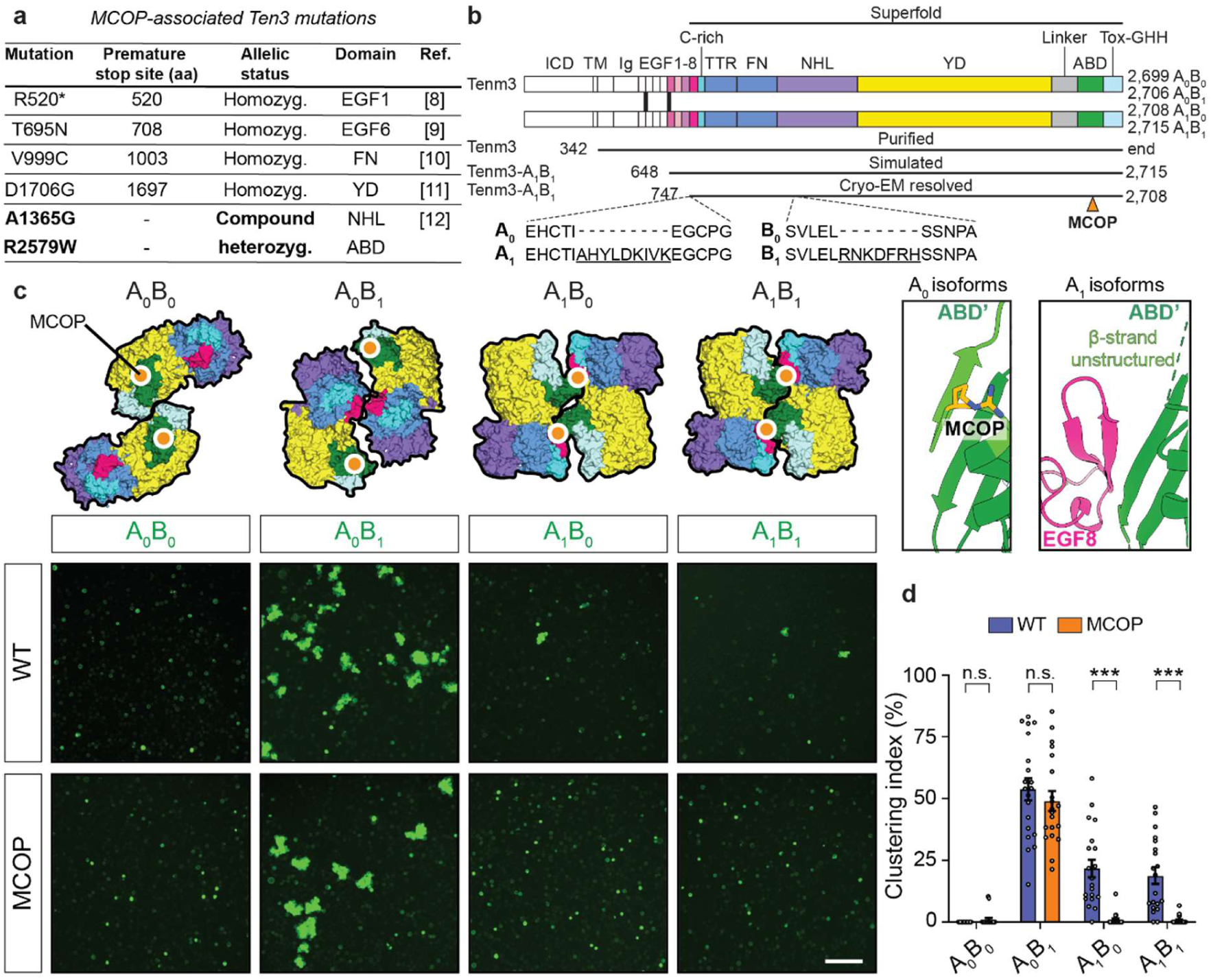
The microphthalmia-associated teneurin-3 (Ten3) mutation is located at the A_1_-specific compact dimer interface and specifically disrupts cell clustering for these isoforms. **a)** Genetic mutations in the human *TENM3* gene that are associated with microphthalmia. Residue numbering corresponds to the A_1_B1 isoform. All mutated residues are identical in the mouse gene, except for T695 (S695 in mouse). In case of frameshifts that lead nonsense mutations, the premature stop codon sites are indicated. Direct nonsense mutations are indicated with *. The allelic status, domain localization, and literature that reported the mutation are also indicated. **b)** Linear representations of the full-length teneurin-3 domain compositions. Splice inserts A and B sequences are underscored and the naming convention for their presence or absence of each splice insert is denoted. Total protein lengths are indicated per isoform, as the splice inserts alter downstream numbering of identical sequences by convention. The rigid superfold (EGF8 through Tox-GHH) is indicated with a black line on top. The black lines below indicate sequence boundaries in this work. ICD, intracellular domain; Ig, immunoglobulin fold; EGF, epidermal growth factor repeat domains 1 through 8; C-rich, cysteine-rich region; TTR, transthyretin-related; FN, fibronectin plug; NHL, NCL, HT2A and Lin-41; YD, tyrosine-aspartate; ABD, antibiotic-binding domain; Tox- GHH, toxin-glycine-histidine-histidine. **c)** Top: Schematic surface representations of the isoform-specific compact dimers^29^. Locations of the MCOP mutation site in each compact dimer is indicated with an orange circle with white outline. EGF repeats that link the subunits are not represented. Right: Insets on the ABD organization for the A_0_ and A_1_ isoforms. To allow EGF8 to bind the ABD domain in the A _1_ isoforms, the outer-most β-strand (light green) of the ABD domain must become unstructured. Bottom: Representative homophilic clustering data of K562 cells expressing the GFP-tagged wildtype (WT) and MCOP mutant full-length Ten3 isoforms. Scale bar is 100 μm. **d)** Quantification of clustering data in panel d. Bar plots represent the percentage that clusters contribute to the total segmented cell area (Clustering index) per image (n=5 images per experiment, N=4 independent experiments, 20 images total). Data are presented as mean ± s.e.m.; ns: not significant, ***: p > 0.001. Specific p-values from left to right are 0.998, 0.638, <0.0001, a<0.0001. Two-Way ANOVA, Šìdák’s multiple comparisons test.

We performed a preliminary assessment of the potential pathogenicity of the R2579W mutation by consulting AlphaMissense^40^. This algorithm predicts mutagenic pathogenicity on basis of structural context, residue conservation across species, genetic variation among humans, and clinical data of mutant-disease association. Although an arginine-to-tryptophan (R-to-W) substitution at position 2579 itself is ambiguous with a pathogenicity score of 0.35 (from 0 to 1), mutating the residues participating in the EGF8-ABD interface is predicted to be highly pathogenic (Supplementary Fig. 1b).

### MCOP mutation disrupts A_1_ homophilic *trans* clustering

To experimentally assess the functional impact of the MCOP mutation on *trans*-cellular interactions, we introduced the R-to-W mutation into N-terminally GFP-tagged full-length mouse Ten3 constructs encoding the four splice variants. These constructs – comprising four wildtype and four MCOP mutant variants - were then expressed in hematopoietic K562 cells to assess their ability to instigate cell clustering through homophilic *trans*-cellular binding (Fig. 1c-d and Supplementary Fig. 1c). As reported earlier^21,29^, cells expressing the wildtype A_0_B_0_ isoform are incapable of forming clusters and remain single cells in suspension. A_0_B_1_ gives rise to the most and largest homophilic clusters, while A_1_B_0_ and A_1_B_1_ consistently conceive less and smaller ones (Fig. 1c). Strikingly, the clustering propensity of the A_1_ isoforms that harbour the microphthalmia mutation is almost completely abolished (Fig. 1c-d). Their total cluster numbers and clustering index are substantially reduced, as are the sizes of the remaining clusters, while both A_0_ clustering phenotypes remained mostly unaffected by the mutation (Fig. 1c-d and Supplementary Fig. 1c). Higher-magnification assessment of the cells reveal no apparent change in GFP fluorescence intensity and display correct localization of all variants to the membrane (Supplementary Fig. 1d). These data show that the MCOP mutation specifically prohibits transcellular clustering of A_1_ isoforms, while having almost no effect on the clustering of A_0_ isoforms.

### Compact dimerization is preserved for all Ten3 MCOP isoforms

To investigate how the MCOP mutation structurally affects the ECD of the four Ten3 splice variants in solution, we performed small-angle X-ray scattering (SAXS) experiments (Fig. 2, Supplementary Fig. 2a-d, and Table 1). For this, we first purified the full ECD proteins of the four splice variants and their mutated MCOP variants using Ni-NTA affinity purification followed by size-exclusion chromatography, both in the presence of calcium (Fig. 1b and Supplementary Fig. 2e). Importantly, no differences in yield or protein quality were observed between the purified protein of the wild-type and mutant variants, again indicating that the MCOP mutation does not affect protein expression.

**Figure 2:**
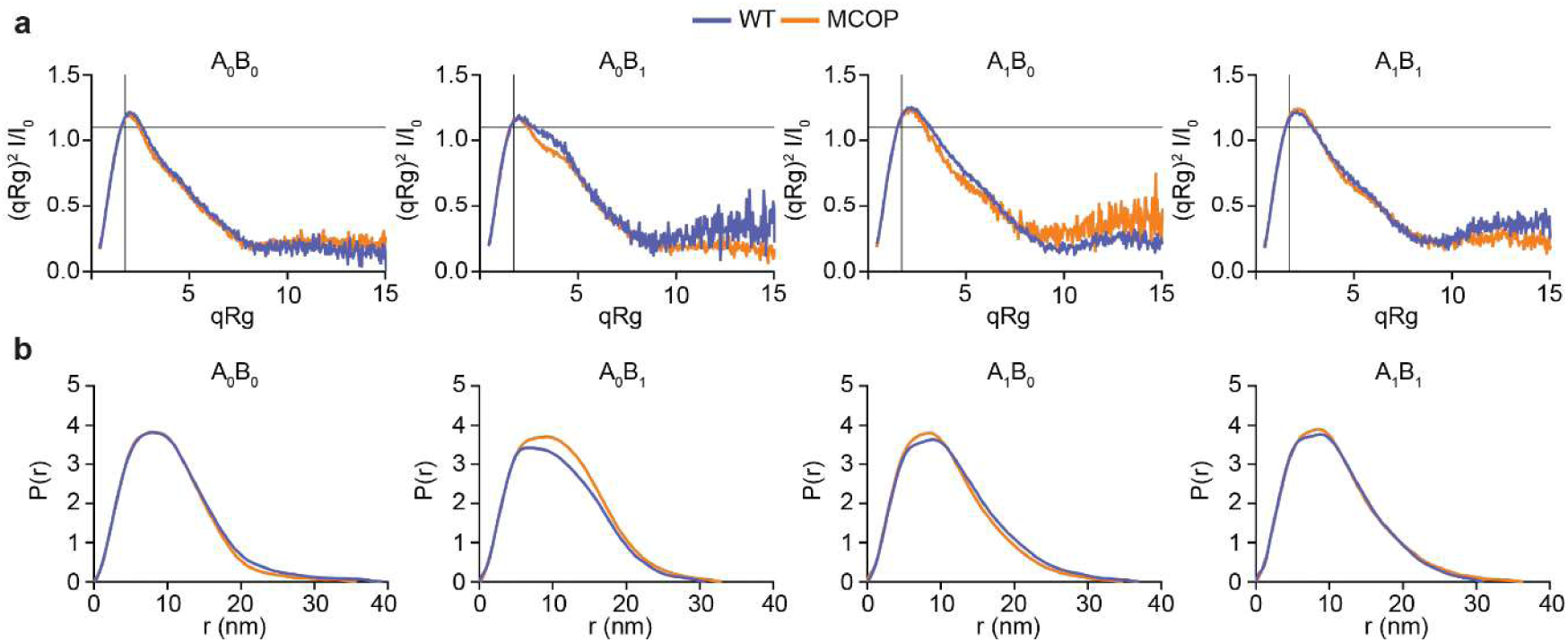
Small-angle X-ray scattering data reveal compact dimer preservation in all teneurin-3 (Ten3) isoforms upon introduction of the MCOP mutation. **a)** Dimensionless Kratky-plots for all Ten3 wildtype (WT, blue) isoforms compared to their respective microphthalmia mutants (MCOP, orange) with maxima of ∼1.1 at *qRg* of √3 (marked with the crossings of the lines), indicating a geometry of a globular particle^41^. **b)** Pair distance distribution functions for all WT isoforms compared to their respective MCOP mutants.

**Table 1:**
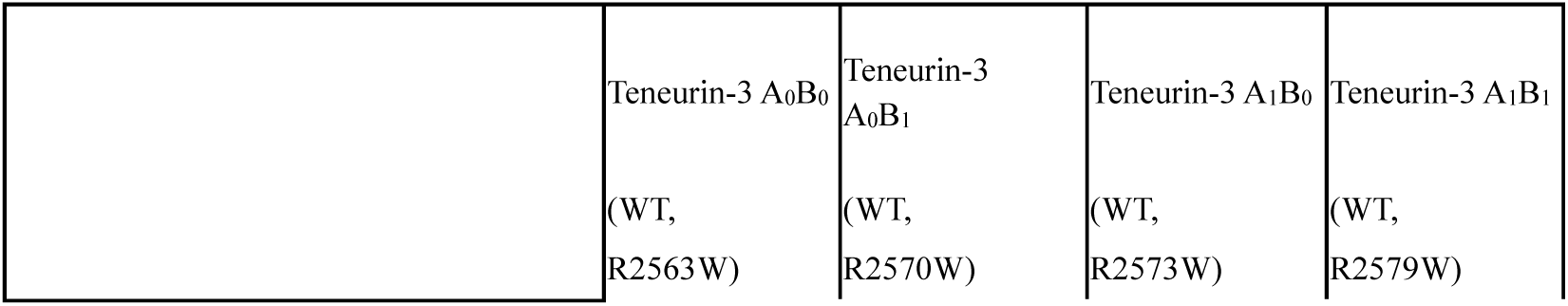
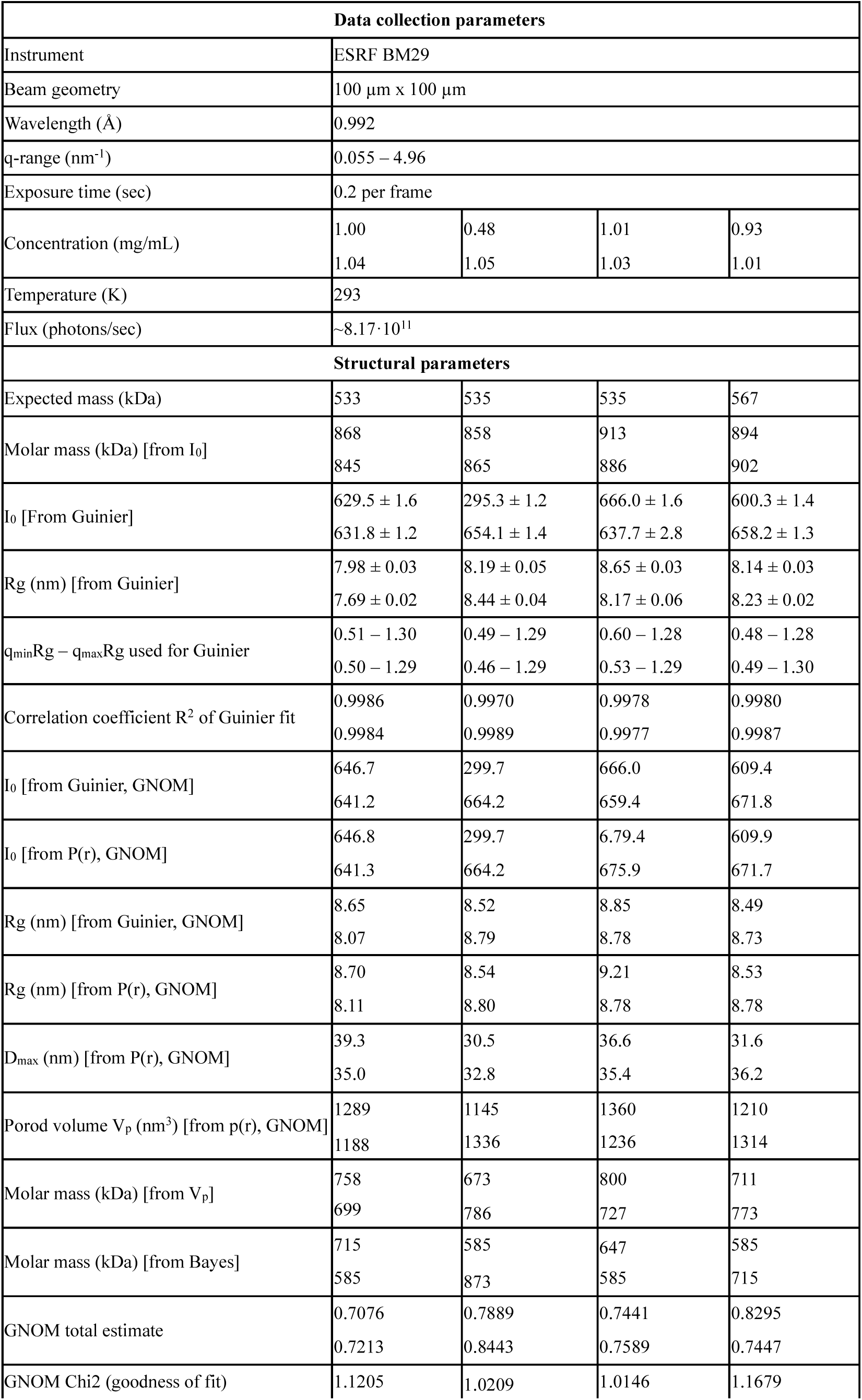
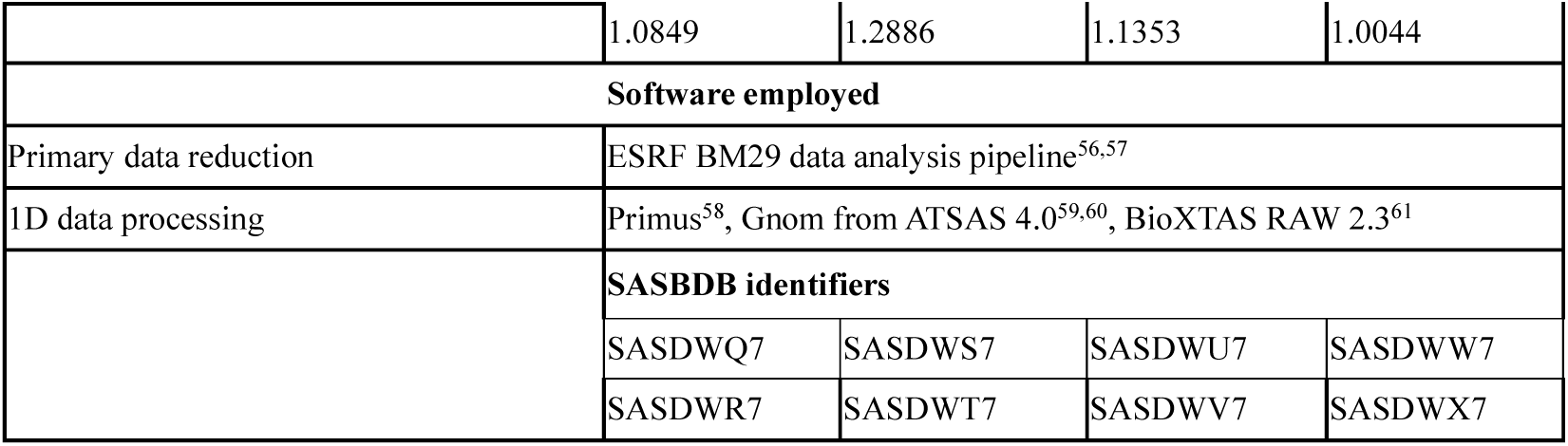
SAXS data collection parameters and analysis.

|  | Teneurin-3 A<sub>0</sub>B<sub>0</sub> | Teneurin-3 A<sub>0</sub>B<sub>1</sub> | Teneurin-3 A<sub>1</sub>B<sub>0</sub> | Teneurin-3 A<sub>1</sub>B<sub>1</sub> |
| --- | --- | --- | --- | --- |
|  | (WT, R2563W) | (WT, R2570W) | (WT, R2573W) | (WT, R2579W) |

| Data collection parameters |  |  |  |  |
| --- | --- | --- | --- | --- |
| Instrument | ESRF BM29 |  |  |  |
| Beam geometry | 100 μm x 100 μm |  |  |  |
| Wavelength (Å) | 0.992 |  |  |  |
| q-range (nm<sup>-1</sup>) | 0.055 – 4.96 |  |  |  |
| Exposure time (sec) | 0.2 per frame |  |  |  |
| Concentration (mg/mL) | 1.00 | 0.48 | 1.01 | 0.93 |
|  | 1.04 | 1.05 | 1.03 | 1.01 |
| Temperature (K) | 293 |  |  |  |
| Flux (photons/sec) | ~8.17·10<sup>11</sup> |  |  |  |
| Structural parameters |  |  |  |  |
| Expected mass (kDa) | 533 | 535 | 535 | 567 |
| Molar mass (kDa) [from I<sub>0</sub>] | 868 | 858 | 913 | 894 |
|  | 845 | 865 | 886 | 902 |
| I<sub>0</sub> [From Guinier] | 629.5 ± 1.6 | 295.3 ± 1.2 | 666.0 ± 1.6 | 600.3 ± 1.4 |
|  | 631.8 ± 1.2 | 654.1 ± 1.4 | 637.7 ± 2.8 | 658.2 ± 1.3 |
| R<sub>g</sub> (nm) [from Guinier] | 7.98 ± 0.03 | 8.19 ± 0.05 | 8.65 ± 0.03 | 8.14 ± 0.03 |
|  | 7.69 ± 0.02 | 8.44 ± 0.04 | 8.17 ± 0.06 | 8.23 ± 0.02 |
| q<sub>min</sub>R<sub>g</sub> – q<sub>max</sub>R<sub>g</sub> used for Guinier | 0.51 – 1.30 | 0.49 – 1.29 | 0.60 – 1.28 | 0.48 – 1.28 |
|  | 0.50 – 1.29 | 0.46 – 1.29 | 0.53 – 1.29 | 0.49 – 1.30 |
| Correlation coefficient R<sup>2</sup> of Guinier fit | 0.9986 | 0.9970 | 0.9978 | 0.9980 |
|  | 0.9984 | 0.9989 | 0.9977 | 0.9987 |
| I<sub>0</sub> [from Guinier, GNOM] | 646.7 | 299.7 | 666.0 | 609.4 |
|  | 641.2 | 664.2 | 659.4 | 671.8 |
| I<sub>0</sub> [from P(r), GNOM] | 646.8 | 299.7 | 6.79.4 | 609.9 |
|  | 641.3 | 664.2 | 675.9 | 671.7 |
| R<sub>g</sub> (nm) [from Guinier, GNOM] | 8.65 | 8.52 | 8.85 | 8.49 |
|  | 8.07 | 8.79 | 8.78 | 8.73 |
| R<sub>g</sub> (nm) [from P(r), GNOM] | 8.70 | 8.54 | 9.21 | 8.53 |
|  | 8.11 | 8.80 | 8.78 | 8.78 |
| D<sub>max</sub> (nm) [from P(r), GNOM] | 39.3 | 30.5 | 36.6 | 31.6 |
|  | 35.0 | 32.8 | 35.4 | 36.2 |
| Porod volume V<sub>p</sub> (nm<sup>3</sup>) [from p(r), GNOM] | 1289 | 1145 | 1360 | 1210 |
|  | 1188 | 1336 | 1236 | 1314 |
| Molar mass (kDa) [from V<sub>p</sub>] | 758 | 673 | 800 | 711 |
|  | 699 | 786 | 727 | 773 |
| Molar mass (kDa) [from Bayes] | 715 | 585 | 647 | 585 |
|  | 585 | 873 | 585 | 715 |
| GNOM total estimate | 0.7076 | 0.7889 | 0.7441 | 0.8295 |
|  | 0.7213 | 0.8443 | 0.7589 | 0.7447 |
| GNOM Chi2 (goodness of fit) | 1.1205 | 1.0209 | 1.0146 | 1.1679 |
|  | 1.0849 | 1.2886 | 1.1353 | 1.0044 |
| **Software employed** |  |  |  |  |
| Primary data reduction | ESRF BM29 data analysis pipeline<sup>56,57</sup> |  |  |  |
| 1D data processing | Primus<sup>58</sup>, Gnom from ATSAS 4.0<sup>59,60</sup>, BioXTAS RAW 2.3<sup>61</sup> |  |  |  |
|  | **SASBDB identifiers** |  |  |  |
|  | SASDWQ7 | SASDWS7 | SASDWU7 | SASDWW7 |
|  | SASDWR7 | SASDWT7 | SASDWV7 | SASDWX7 |

It is directly apparent from the similarity between wildtype and mutant scattering curves that the mutations do not disrupt the overall compact dimeric organization of any isoform (Fig. 2 and Supplementary Fig. 2a-d). Moreover, it was previously shown by SAXS that calcium chelation provokes a significant extension of the overall structure for all isoforms^29^. We do not observe such a decompaction in the mutant dataset. While the SAXS curves reveal no major change in the global architecture of the mutant proteins, some subtle differences between the wildtype and mutant isoforms can be observed. Firstly, the A_1_B_0_^MCOP^ and A_1_B_1_^MCOP^ dimensionless Kratky plots exhibit steeper peak decay compared to their wildtype counterparts (Fig. 2a), which indicates a more compact core structure of the mutant proteins^41^. This can also be seen in the pair distance distribution plots (P(r)) with more symmetrical peaks compared to the wildtype (Fig. 2b). While A B ^MCOP^ has higher radius of gyration (R_g_) and maximum particle dimension (D_max_) values than its wildtype (Table 1), the sharper and more symmetrical peaks in the Kratky plot and P(r), respectively, indicate that this may reflect increased flexibility of its peripheral domains.

Furthermore, while not showing a large effect in the clustering assays, the pathogenic mutation also has an effect on the in-solution A_0_B_1_ structure. The R_g_ and D_max_ values indicate a less compact A B ^MCOP^ ECD compared to its wildtype counterpart (Fig. 2b and Table1). The sharper peak in the Kratky plot and increased symmetry in P(r) again indicate a more compact core with other intramolecular interactions possibly loosening up. The MCOP substitution has the least effect on the structure of the A_0_B_0_ isoform. Altogether, the SAXS data demonstrate that the Ten3 compact *cis* dimers do not dissociate in the presence of the MCOP mutation, and that the A_1_ isoform adopt more compact core structures.

### Molecular dynamics simulations reveal A_1_^MCOP^ compact dimer stabilization

Since the SAXS curves reveal preservation of the overall protein architecture of the A_1_ mutant ECDs, local changes in the mutant versus wildtype structure may explain the observed cellular effects. To computationally probe the effect of the R-to-W MCOP substitution at high spatiotemporal resolution, we employed molecular dynamics (MD) simulations, focusing on the splice insert A-containing isoforms. We used the A_1_B_1_ compact dimer structure (PDB: 8R50) as the representative A_1_ starting structure, with four important features. First, EGF repeats 5, 6 and 7 were simulated to include the natural covalent link of the two subunits (Fig. 1b and Supplementary Fig. 3a). Second, glycan chains were not included in the simulations. Third, on basis of sequence conservation^34^, six structural calcium ions were placed in the two C-rich domain to stabilize the complex. Fourth, the loop and β-strand that were originally missing from the cryo-EM density were built into the structure to assess the effect of the R2579W mutation on these residues (Supplementary Fig. 3b). Three repeats of 1-μs simulations were run for the wildtype and the R2579W MCOP mutant.

Upon analysis of all interactions between the two subunits in each compact dimer, we find comparable numbers of hydrogen bonds in wildtype and mutant (Fig. 3a) but an increased amount of ionic bonds in the wildtype (Fig. 3b). We next looked at direct interactions formed by the wildtype arginine and mutant tryptophan at position 2579. While both interact a similar proportion of the time with the more N-terminal EGF repeats, the arginine interacts much more with the YD shell (Fig. 3c). Closer inspection reveals inter-subunit salt bridges formed exclusively by the wildtype arginine with D1647 in the YD shell (Fig. 3d) and with E718 in EGF7, thereby accounting for the increased number of ionic bonds in the wildtype. Residues Y743, L744, I747 in splice insert A and W769 in EGF8 form a hydrophobic pocket in one subunit that interacts with the mutant R2579W substitution of the other subunit (Fig. 3e). Splice insert A was shown to be pivotal for the formation of the A_1_-specific compact dimers^29^. Here, it also directly interacts with the MCOP-associated R-to-W substitution.

**Figure 3:**
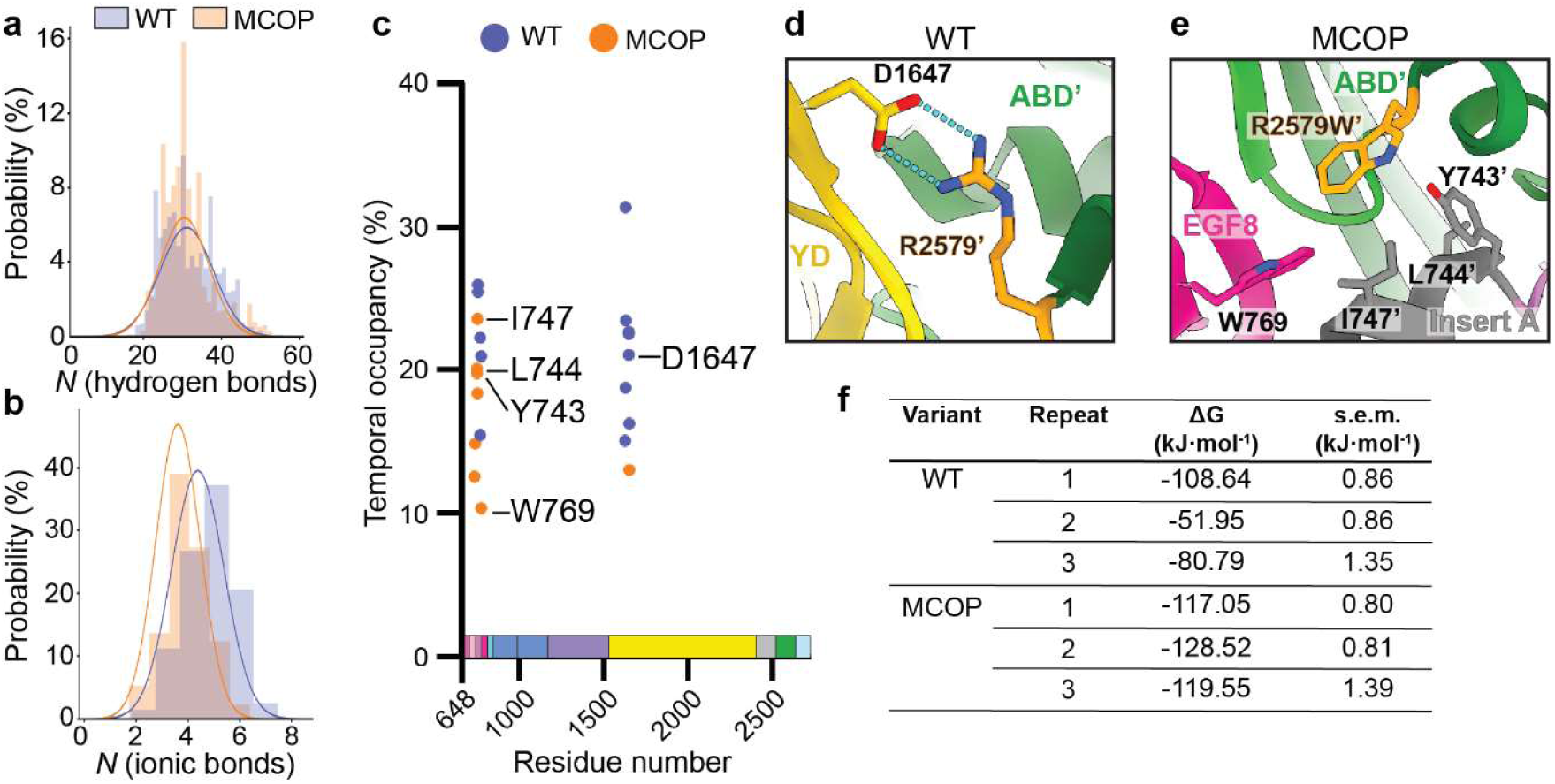
Molecular dynamics simulations predict MCOP compact dimer stabilization by local differences in interactions with the wildtype vs. mutated residue. **a)** Probability distribution of hydrogen bonds formed between subunits in the compact dimer throughout the simulation for wildtype (WT) versus the MCOP mutant. **b)** Probability distribution of ionic bonds formed between subunits in the compact dimer throughout the simulation for WT versus the MCOP mutant. Bell curves in panels a-b are Gaussian fits to the histograms. **c)** Proportions of the time spent (Temporal occupancy) by WT arginine (blue) vs. MCOP tryptophan (orange) within 4 Å of any residue on the other subunit of the compact dimer. Indicated residues are the high-occupancy partners depicted in panel d-e. **d)** Inset of the ionic bond formed between WT arginine 2579 on one subunit and aspartate 1647 in de YD shell of the other subunit. **e)** Inset of the hydrophobic region spanned by tyrosine 743, leucine 744, isoleucine 747, and tryptophan 769 on one subunit that directly interacts with the MCOP tryptophan 2579 substitution on the other subunit. Domain colour codes of panels c-e correspond to the description in Fig. 1b. **f)** Total binding free energy (ΔG) of the compact dimer interface in each 1-μs simulation. Values represent mean and s.e.m. of binding affinities calculated from the last 500 ns of each trajectory with 1 ns between each frame. **g)** Probability distribution comparison of numbers of hydrogen bonds and numbers of ionic bonds between subunits throughout WT and MCOP simulations. Apostrophes (’) in panels d, and e are used to distinguish between residues located on one subunit in the compact dimer vs. localization on the other subunit.

On a more global scale, a decrease in R_g_ and r.m.s.d. of the mutant compact dimer is observed toward the end of the simulations (Supplementary Fig. 3c-g). Crucially, the MCOP mutation gives rise to a stabilized compact dimer, as characterized by a more negative binding free energy (Δ*G*) of –121.71 ± 3.48 kcal ⸱ mol^-1^ (mean ± s.e.m. between 3 simulations), compared to –80.46 ± 16.37 kcal ⸱ mol^-1^ for the wildtype (Fig. 3f). Given the higher number of ionic bonds in the wildtype, the more stable binding energy of the mutant most likely results from an increased contribution by hydrophobic interactions. It should be noted that the loop and β-strand remain structured throughout all simulations (data not shown). Thus, the mutation itself does not affect the loop and β-strand in hindering the EGF8-ABD interaction.

The simulations also revealed an unexpected feature of the Ten3 subunits. Throughout all wildtype and MCOP simulations, calcium ions were bound to a negatively charged patch on the YD shell near the ABD domain (Supplementary Fig. 3h). Other, more transient binding sites were also observed at the EGF5-EGF5 interface and at the NHL/TTR domain (data not shown). Part of the YD patch corresponds to the EGF6-YD/ABD contact previously described for the A_0_ isoforms and may thus play a role in A_0_ compact dimer formation^29^. Of the seven negatively charged amino acids in this region, four are conserved across all human and mouse teneurin family members (Supplementary Fig. 3i). The remaining three residues are altered or missing in mouse Ten1 (only two in human Ten1), potentially having implications for the calcium-dependence of mouse and human Ten1.

### Biophysical validation of A_1_-specific stabilization by the MCOP mutation

The MD simulations suggest increased stability in the A_1_ compact dimer conformations, prompting us to test this with biophysical approaches. We first assessed the overall stability of the purified Ten3 wildtype and mutant ECDs by performing a thermal stability assay (TSA). A significant increase in melting temperature (T_m_) is only observed for the A_1_ isoforms (Fig. 4a-b); The average T_m_ is increased by 0.65 and 0.43 °C for the A_1_B_0_ and A_1_B_1_ isoforms, respectively. These A_1_-specific differences were lost in the absence of calcium (Supplementary Fig. 4a) - when the compact dimer conformation is disrupted^29^, indicating that the MCOP mutation specifically affects their compact dimer interface.

**Figure 4:**
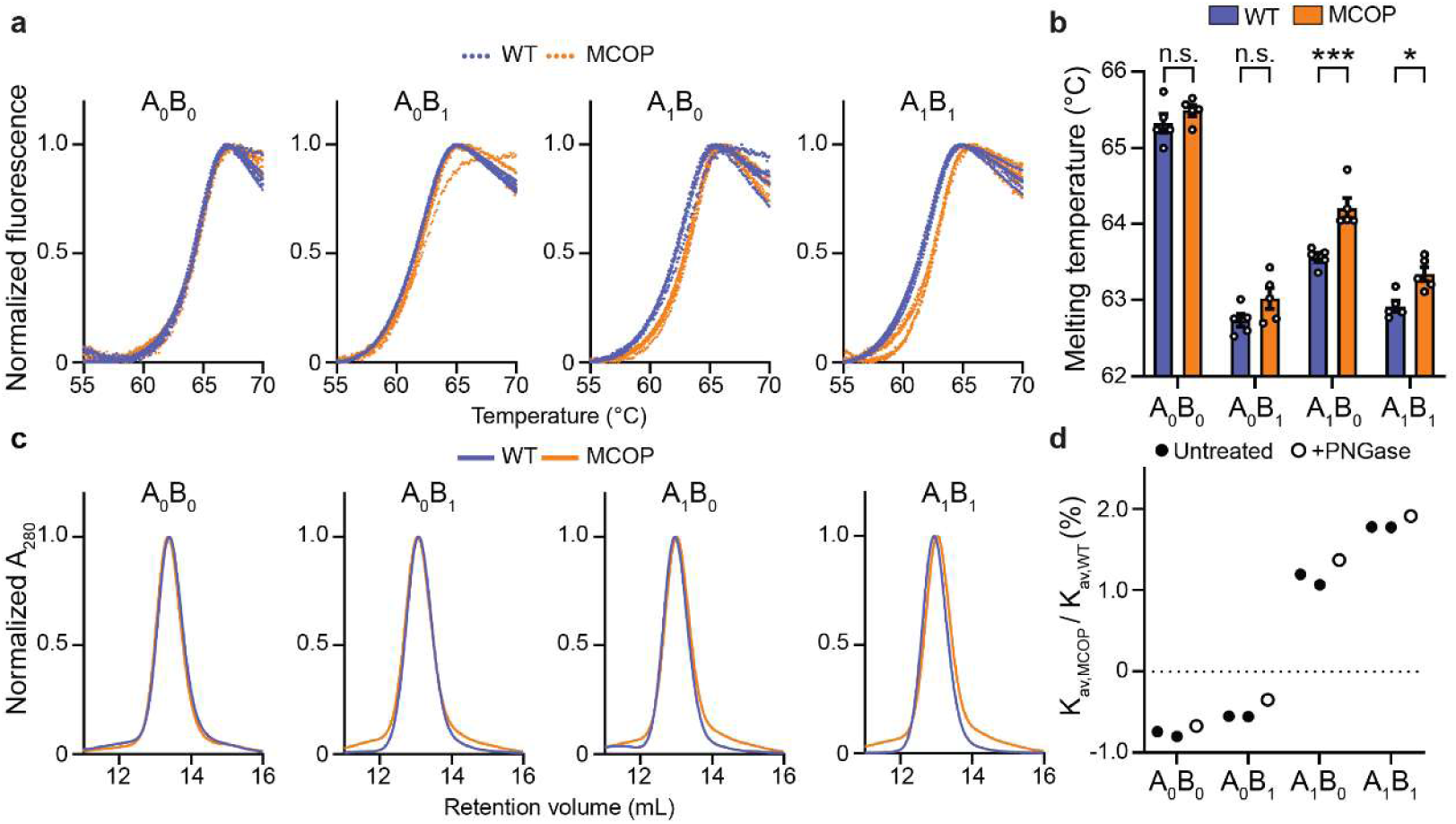
Biophysical validation of A_1_-specific compact dimer stabilization by the MCOP mutation. **a)** Thermal stability assay melting curves of all Ten3 wildtype (WT, blue) isoforms compared to their respective microphthalmia mutants (MCOP, orange). Five technical repeats per variant are plotted. **b)** Melting temperatures derived from the melting curves in panel a. Data are presented as mean ± s.e.m.; ns: not significant, *: p > 0.05, ***: p > 0.001. Specific p-values from left to right are 0.658, 0.178, 0.0003, and 0.019. Two-Way ANOVA, Šìdák’s multiple comparisons test. **c)** Analytical size-exclusion chromatography (aSEC) curves of WT isoforms compared to their respective MCOP mutants. **d)** Quantification of MCOP-mutation induced peak shifts in aSEC curves from panel c. The partition coefficient of each mutant (K_av,MCOP_) run was divided by the partition coefficient of the wildtype (K_av,WT_) run in the same experiment. Single-repeat experiments are also plotted where all Ten3 wildtypes and mutants were treated with PNGase (+PNGase) to strip the surface-exposed glycans.

We next performed analytical size-exclusion chromatography (aSEC) to probe whether the compact dimer conformations were stabilized on population level. As shown in figure 4c, the A_1_^MCOP^ variants consistently elute later than their respective wildtypes, while A_0_ mutants reproducibly elute slightly earlier than theirs. We determined the relative change in aSEC partition coefficient (K_av_) between each isoform’s wildtype and respective MCOP mutant (K_av,Mut_ / K_av,WT_) to quantify the peak shift and compare between the experimental repeats (Fig. 4d).

We were also interested to test whether the peak shifts could emerge from changes in molecular weights due to altered glycosylation profiles (17 predicted glycosylation sites on Ten3 ECD). All purified Ten3 wildtype and their MCOP isoforms were digested with PNGase to remove their entire glycan chains (Supplementary Fig. 4b). Although all variants eluted later due to the stripping of the glycans, the relative changes within each isoform’s wildtype-mutant pair were preserved (Fig. 4d). This indicates that the observed shifts in aSEC partition coefficients are not likely the result of a change in mass due to altered glycan profiles but instead result from stabilized A_1_ compact dimers.

### MCOP mutation restricts conformational freedom of the A_1_ compact dimer

To obtain high-resolution experimental data of the pathogenic compact dimer, we performed cryo-EM single particle analysis (SPA) on the full-length extracellular region of the Ten3 A_1_B_1_^MCOP^ mutant (Fig. 1b, 5a, Supplementary Fig. 5a-d, and Table 2). Similar to the wildtype dataset ^29^, 3D classes representing so-called ‘open’ and ‘closed’ conformations – where the closed conformation has an additional YD-YD contact – are also observed for the mutant (Supplementary Fig. 5d-e). SPA of the particles in closed conformation resulted in an overall resolution of 3.0 Å of the compact dimer conformation (Fig. 5a, Supplementary Fig. 5d, and Table 2). The mutant compact dimer is highly comparable to the previously published wildtype compact dimer of Ten3 A_1_B_1_, with an r.m.s.d. of 0.4 Å over 3567 Cα’s (Fig. 5b). R2579 was not resolved in the wildtype compact dimer (Fig. 5c)^29^, because, together with the β-strand directly flanking it, it was found to be displaced to enable EGF8-ABD interaction. While the loop and β-strand are still largely unstructured in the R2579W mutant, extra density is now observed that represents the tryptophan flanking this unstructured region (Fig. 5c). This extra density plugs into a hydrophobic pocket formed by residues L764 and W769 on EGF8, and L2575, L2578, V2595 on the ABD (Fig. 5c-e). These amino acids participate in the EGF8-ABD extended β-sheet. Additionally, as predicted by the MD simulations, residue I747 from splice insert A is also resolved and may participate in this hydrophobic interaction as well (Fig. 3e and Fig. 5c-e).

**Figure 5:**
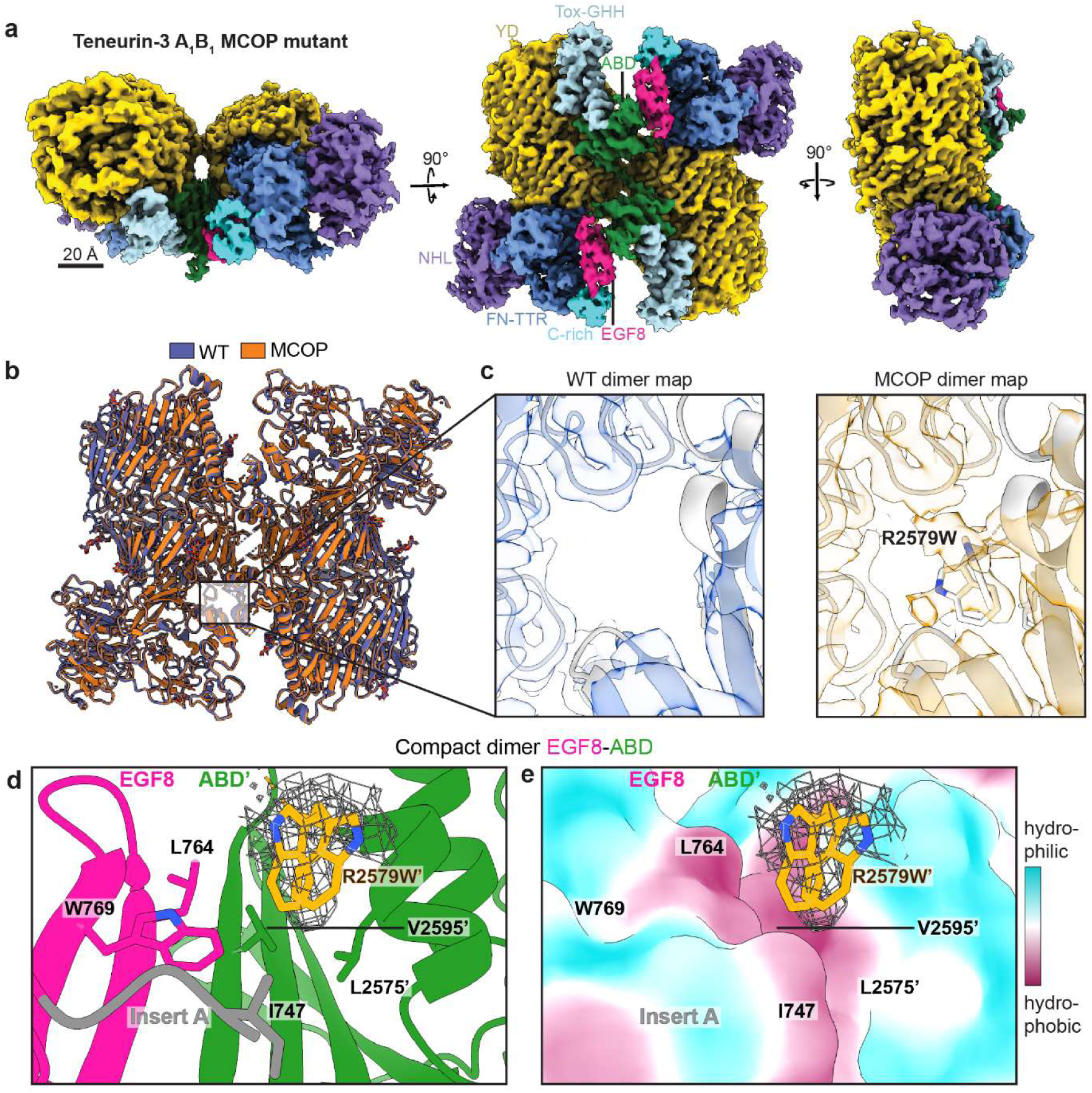
High resolution cryo-EM structure of the Ten3-A_1_B_1_ R2579W (MCOP) compact dimer. **a)** Density map of the Ten3-A_1_B_1_ R2579W (MCOP) compact dimer at 3.0 Å resolution at three orthogonal views (EMD-52854). Scale bar is 20 Å. **b)** Overlay of the wildtype (WT, PDB:8R50) with the MCOP (PDB:9IGS) compact dimer cryo-EM structures. **c)** Close-up of the compact dimer interface showing differences in electron density between WT and MCOP at residue position 2579. The MCOP reconstruction shows extra density at the location of R2579W. The structures are coloured white and the density maps in purple and orange transparency. **d)** Close up of the MCOP EGF8-ABD contact, including two rotamers of R2579W and isoleucine 747 in splice insert A. Density within 2 Å of R2579W is shown as mesh. Apostrophes (’) are used to distinguish between residues located on one subunit in the compact dimer vs. on the other subunit. **e)** Hydrophobic surface representation of the view shown in panel c, with the hydrophobicity colour coding legend is shown on the right.

**Table 2:**
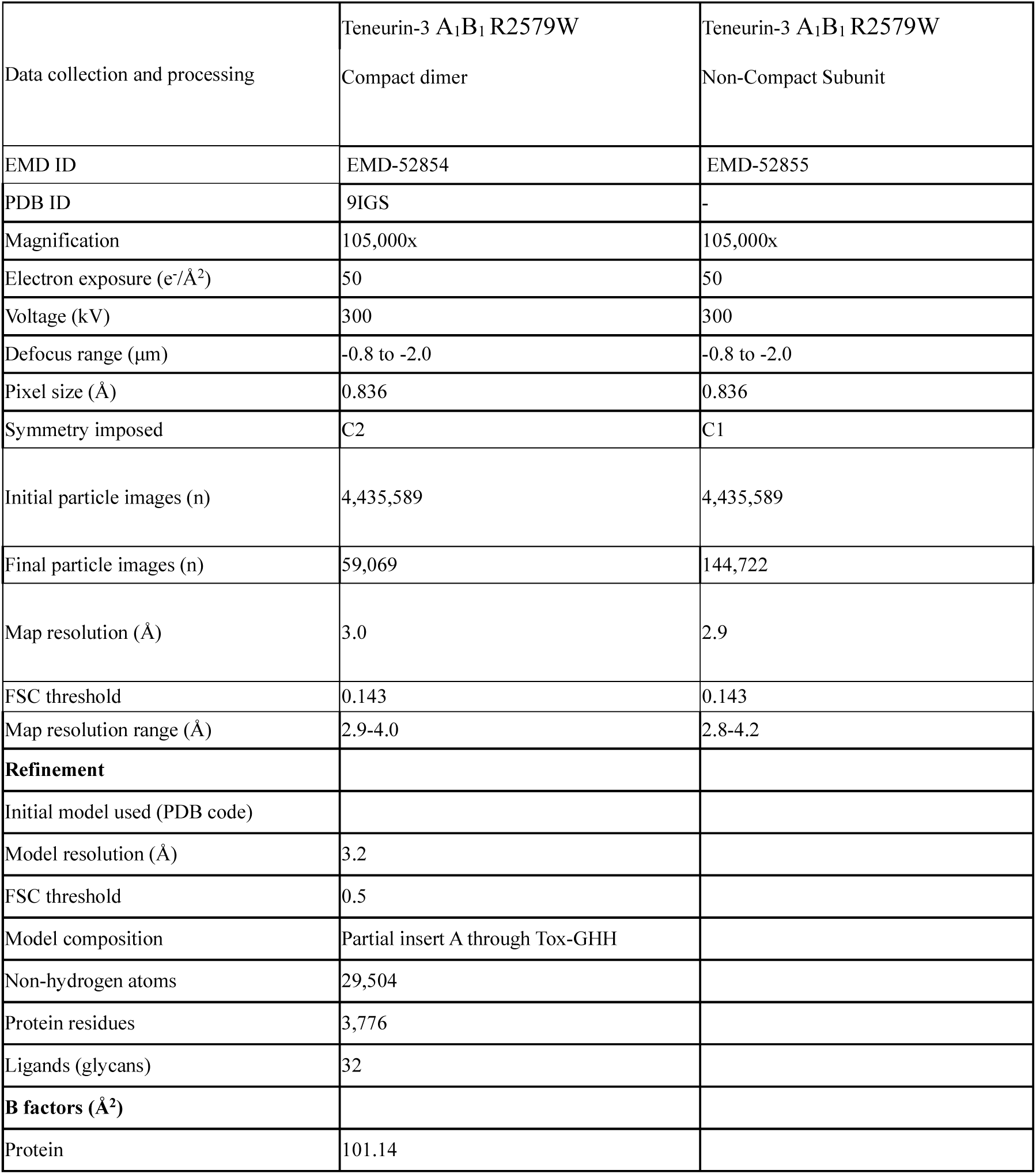
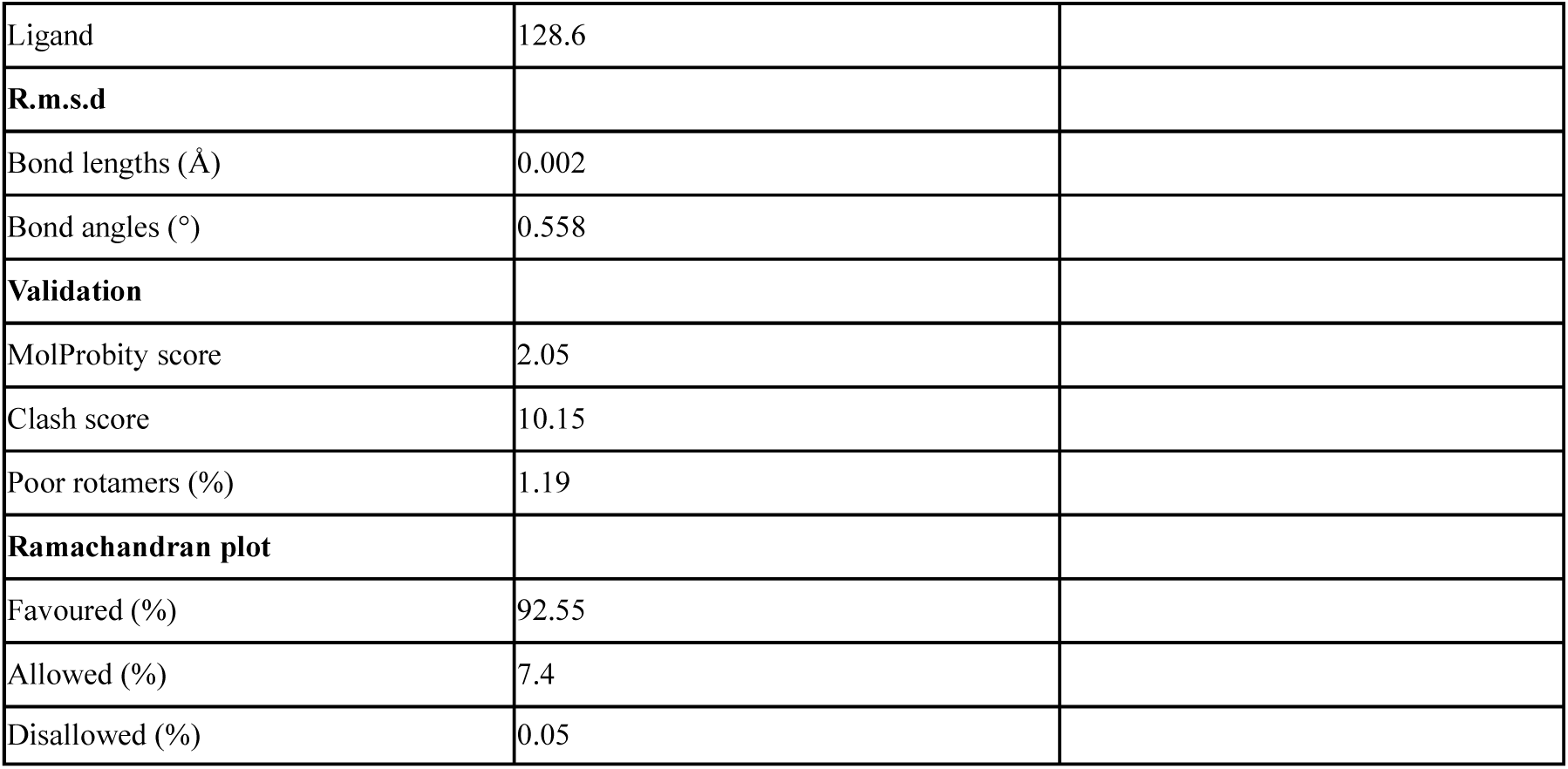
Cryo-EM data collection and refinement statistics.

|  |  |  |
| --- | --- | --- |
| Data collection and processing | Teneurin-3 A<sub>1</sub>B<sub>1</sub> R2579W<br>Compact dimer | Teneurin-3 A<sub>1</sub>B<sub>1</sub> R2579W<br>Non-Compact Subunit |
| EMD ID | EMD-52854 | EMD-52855 |
| PDB ID | 9IGS | - |
| Magnification | 105,000x | 105,000x |
| Electron exposure (e<sup>-</sup>/Å<sup>2</sup>) | 50 | 50 |
| Voltage (kV) | 300 | 300 |
| Defocus range (μm) | -0.8 to -2.0 | -0.8 to -2.0 |
| Pixel size (Å) | 0.836 | 0.836 |
| Symmetry imposed | C2 | C1 |
| Initial particle images (n) | 4,435,589 | 4,435,589 |
| Final particle images (n) | 59,069 | 144,722 |
| Map resolution (Å) | 3.0 | 2.9 |
| FSC threshold | 0.143 | 0.143 |
| Map resolution range (Å) | 2.9-4.0 | 2.8-4.2 |
| **Refinement** |  |  |
| Initial model used (PDB code) |  |  |
| Model resolution (Å) | 3.2 |  |
| FSC threshold | 0.5 |  |
| Model composition | Partial insert A through Tox-GHH |  |
| Non-hydrogen atoms | 29,504 |  |
| Protein residues | 3,776 |  |
| Ligands (glycans) | 32 |  |
| **B factors (Å<sup>2</sup>)** |  |  |
| Protein | 101.14 |  |

|  |  |
| --- | --- |
| Ligand | 128.6 |
| **R.m.s.d** |  |
| Bond lengths (Å) | 0.002 |
| Bond angles (°) | 0.558 |
| **Validation** |  |
| MolProbity score | 2.05 |
| Clash score | 10.15 |
| Poor rotamers (%) | 1.19 |
| **Ramachandran plot** |  |
| Favoured (%) | 92.55 |
| Allowed (%) | 7.4 |
| Disallowed (%) | 0.05 |

The mutant A_1_B_1_ non-compact particle was also reconstructed to evaluate the organization of the ABD domain when it is not in contact with EGF8 (Supplementary Fig. 5d-e and Table 2). The β-strand and the loop that harbours the R2579W mutation are not resolved anymore in the non-compact particles, while they were resolved in the wildtype (Fig. 5e). This indicates that this loop is already destabilized in non-compact subunits and may thereby also facilitate compact dimer formation by not obstructing the EGF8-ABD interaction.

Note that of all picked particles from the MCOP dataset, roughly 10% were classified as compact dimers, compared to the ∼7% of compact dimers in the previous wildtype dataset (434,882 dimers out of 4,435,589 particles total for the mutant vs. 159,924/2,196,963 for the wildtype^29^). This slight increase in compact dimeric particles may also reflect the stabilizing effect of the MCOP mutation.

We further analysed the wildtype and mutant heterogeneous particle datasets to determine the principal modes of movement (PMMs) of the compact dimers (Supplementary Fig. 5d). Notably, the first three PMMs found in the experimental wildtype and mutant cryo-EM datasets are highly comparable to those calculated from the MD simulations (and Supplementary Movies 1-3). For the most dominant experimental PMM, in which the two subunits transition between the open and closed conformations in a so-called ‘butterfly movement’ (Supplementary Fig. 6 and Supplementary Movie 1), the wildtype opens more widely compared to the mutant. We quantified this by rigid body fitting single subunits into the extremes of the respective PPM and determining the r.m.s.d. between the resulting complexes. The open-to-closed r.m.s.d. of the wildtype and MCOP mutant are 4.7 and 4.4 Å, respectively (Supplementary Fig. 6a-b). The further opening of the wildtype becomes more apparent when comparing wildtype open state with that of the mutant at an r.m.s.d. of 1.8 Å (Supplementary Fig. 6c), while the closed states only differ by 0.9 Å (Supplementary Fig. 6d). This analysis thus indicates a restricted degree of inter-subunit conformational freedom in the MCOP compact dimer.

The findings from the cryo-EM structure align well with the MD simulations regarding the direct interactions of the mutant tryptophan, and with SAXS regarding the more compact core structure of the Ten3 A_1_ ECD.

## Discussion

Neurons use physical interactions between cell adhesion molecules at the cell surface as guidance cues to realize proper integration into neuronal networks^12–14,16,22,29,33,42,43^. Disruption of transcellular protein-protein interactions can lead to miswiring of the nervous system with clinical implications, referred to as connectopathies^44,45^. In this work, we deliberate on various Ten3 mutations associated with microphthalmia (MCOP) and focus on the C-terminal arginine-to-tryptophan mutation (R2579W in the A_1_B_1_ isoform). We previously shed light on large conformational differences instigated by the splice inserts A and B, located in between EGF7-8 and in the NHL domain, respectively ^29^. Aside from the constitutive disulfide linkage of two Ten3 monomers at the flexible EGF membrane tethers, the monomers in all four isoforms form additional non-covalent contacts, which we refer to as compact dimer interfaces. Here, we structurally mapped the MCOP missense mutation to the intramolecular compact dimer interface shared exclusively by the two isoforms that contain splice insert A (A_1_B_0_ and A_1_B_1_). The mutation leads to disruption of transcellular adhesion specifically in A_1_ isoforms. Small-angle X-ray scattering (SAXS) data reveal a preservation of all compact conformations, with MD simulations and biophysical assays showing a stabilization of this compactness. Using high-resolution cryo-EM, we resolve a novel interaction of the pathogenic residue on one subunit with a hydrophobic pocket on the other. This interaction and the resulting stabilization manifests as reduced conformational freedom of the mutant compact dimer. This agrees well with the stronger overall compaction of the mutant structures compared to their wildtypes observed in SAXS analysis.

Together, our data strongly suggest that conformational and energetic stabilization of the compact dimers in *cis* is causally linked to the isoform-specific inhibition of cell-cell adhesion in *trans*. We propose that stabilization in *cis* could prevent dimer-of-dimer interactions between cells by two primary routes (Fig. 6 and Supplementary Fig. 7). Firstly, the dimer conformation forms a yet unresolved dimer-of-dimer contact with the adjacent cell, where the formation of this contact requires some conformational flexibility of the compact *cis*-dimer (Supplementary Fig. 7a-b). The restriction of conformational freedom in *cis* – as found in our mutant cryo-EM dataset - would then prevent the adoption of a conformation that is compatible with dimer-of-dimerization. Alternatively, wildtype *trans*-adhesion might require complete dissociation of the compact *cis* dimer before formation of two *trans* compact dimer interfaces between the four subunits of two covalent dimers (Supplementary Fig. 7a-b).

**Figure 6:**
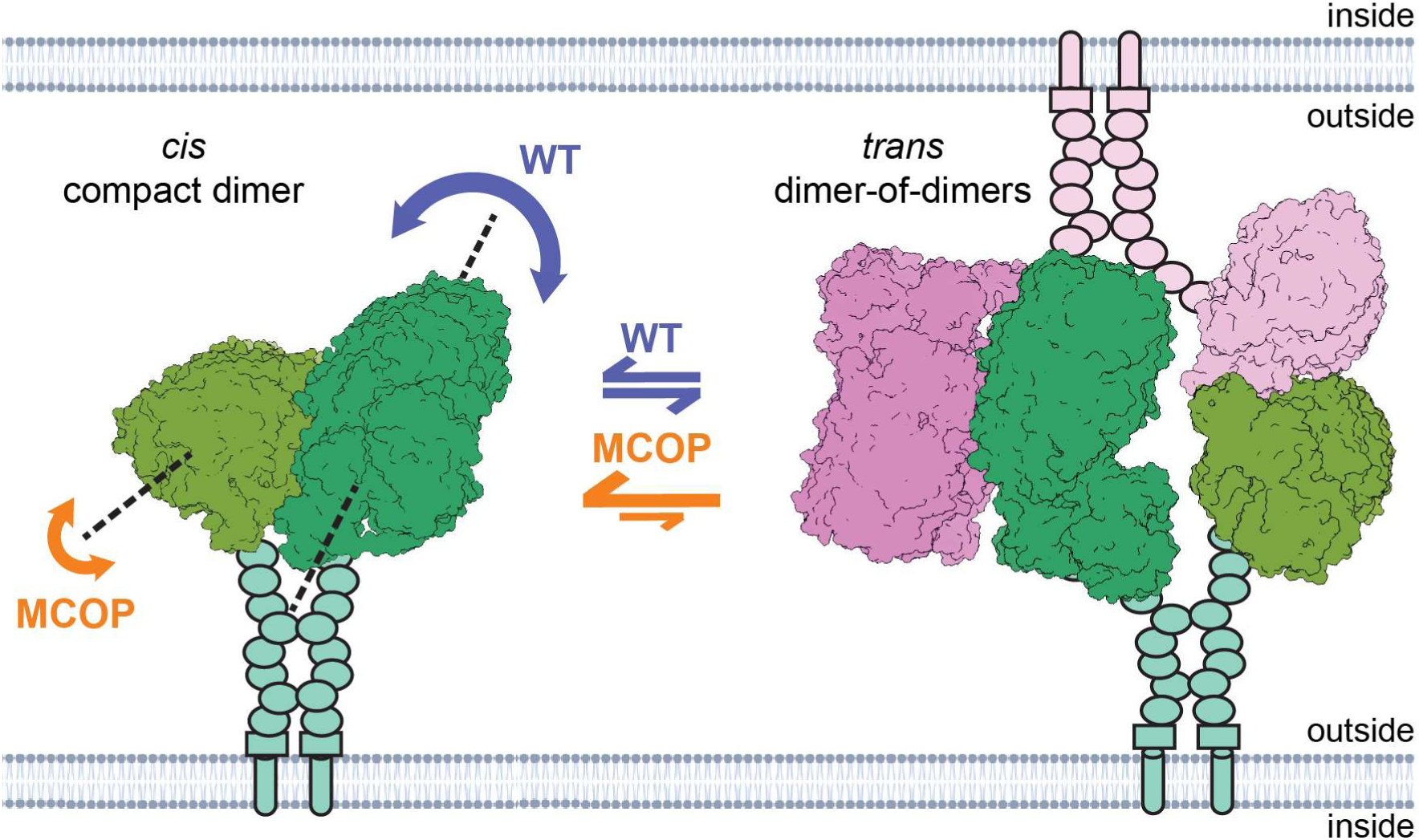
Model for MCOP-induced inhibition of *trans* adhesion via stabilization of the *cis* compact dimer. Schematic visualization of a Ten3 A_1_ *cis* compact dimer (green) with more conformational freedom between the two subunits in the wildtype (WT) case compared to the stabilized MCOP mutant. We hypothesize that this conformational freedom or complete dissociation of the *cis* compact dimer is required for *trans* interactions with another covalent dimer (pink); in the presence of the mutation, each subunit is more likely to undergo a compact dimer interaction in *cis* than in *trans*. All Ten3 molecules are visualised as attached to the extracellular (outside) side of the membrane via a transmembrane helix (tube), an Ig domain (rectangle), and eight EGF repeats (ovals) in the same colour as the attached subunits. Intracellular (inside) domains are omitted for clarity.

While homophilic transcellular adhesion is completely lost for the A_1_ isoforms in our clustering assay, a less dominant yet statistically significant reduction is also found in cluster size and count for A_0_B_1_ (Supplementary Fig. 1c). SAXS data in addition reveal a mutagenic effect on the compact structure of this isoform (Fig. 2). Given the reverse peak shift of A B ^MCOP^ in aSEC (Fig. 4c-d), we hypothesize that the MCOP mutation in A_0_B_1_ does not stabilize the compact dimer interface directly, as is the case for the A_1_ variants, but may be explained by an increased flexibility of smaller or more peripheral domains.

Direct heterophilic *trans* interactions between teneurins and adhesion GPCR latrophilin (Lphn) are crucial for synapse organization and act as repulsive guidance cues for outgrowing neurons^21,22,28,32,33,38,46–48^. From the model presented here, it may be expected that the increased stability of Ten3 A_1_ isoforms will not detrimentally affect the *trans* interaction with Lphn, as the MCOP mutation is not located near the Lphn binding site on the YD shell in any of the isoforms’ compact dimeric organizations^29,32,33^. We also do not find indirect shielding of the Lphn binding sites through MCOP- induced structural reorganizations in Ten3. Within this context, it is worth noting that studies into the Ten-Lphn interaction primarily concern wiring of the hippocampus^22,24,46,47^, while studies that focus on teneurins alone report effects in the optic system and vision^19,23,25^. Perhaps teneurin homophilic interactions, which we show here to be selectively disrupted by the MCOP mutation, are more specifically involved in wiring the visual system.

Missense mutations have been described previously to affect alternative splicing resulting in a pathogenic outcome. For example, in frontotemporal dementia and Alzheimer’s disease, various mutations in *MAPT* gene result in imbalanced production of the native 3R and 4R splice isoforms of tau protein^49^. Splicing mutations that give rise to unnatural proteoforms have also been reported by, for instance, skipping canonical exons or generating new ones^50,51^. A missense mutation within an alternatively spliced exon has been described for the Noonan syndrome, a developmental disorder. Here, the mutation affects only KRAS isoform B but not A, as exon 6 – where the mutation occurs – is exclusive to this isoform^52^. Finally, a G12D mutation is present in an exon shared between KRAS4A and -B isoforms that differentially affects protein-protein interactions of these isoforms^53^. To our best knowledge, no missense mutations are described that are expressed in all isoforms but only affect the function of some, as is the case for R2579W in Ten3. This can be explained by the unique mechanism where its splicing isoforms alternatively hide and expose binding motifs through substantial molecular reorganization of the ECD^54^.

The unique patient-derived R-to-W point mutation we investigate here allows us to study the dysfunctional molecule *in vitro*, whereas the other nonsense truncation (Fig. 1a) would only permit knockdown approaches in more complex biological context. Moreover, its isoform-specific knockdown phenotype paves a way to explore the roles of the existing splicing isoforms. Although the expression of different teneurin family members has recently been chartered throughout mouse development^24^, and the differential expression of isoforms in mouse hippocampus has been demonstrated^21^, isoform- specific roles in the brain remain unknown. Getting high-resolution insight into the compact dimerization of the A_0_ variants may provide the design of similar isoform-specific knockdown mutations to complement this genetic toolbox for investigating the biological roles of Ten3 alternative splicing.

Ten3 is crucial for axonal pathfinding and neuronal integration in different parts of the brain across species^17–19,21–25^. Loss of either homophilic or heterophilic interactions with outgrowing Ten3- expressing axons leads to global miswiring in mouse hippocampus^22^. A more recent study on fly Ten- m revealed that the effective level of teneurin on pre- and postsynapses need to match for the contact to be established^55^. Disruption of any of these sensitive balances in partner matching may greatly impact neuronal wiring. Given the predominant expression of Ten3 in the hippocampus and visual system, the mutagenic alteration of teneurin homophilic *trans*-cellular adhesion presented here may very well contribute to the neurological and visual defects observed in MCOP. Our findings provide the first molecular evidence for the pathogenicity of MCOP-associated R-to-W substitution by isoform-specifically interfering with Ten3 *trans*-cellular recognition. This may unsettle the repulsive and attractive cues of teneurin-expressing neurons that are required for proper synaptic matching and could thereby lead to neurological impediments and ocular malformations found in MCOP patients.

## Methods and material

### Statistics and reproducibility

No statistical method was used to predetermine sample size. No data were excluded from the analyses. The experiments were not randomized, and the investigators were not blinded to allocation during experiments and outcome assessment.

### Cloning

Plasmids encoding the his-tagged extracellular domains and GFP-tagged full-length wildtype teneurin- 3 (Ten3) isoforms were generated as described previously^29^. The constructs containing the arginine-to-tryptophan substitution (R2563W, R2570W, R2572W, and R2579W in A_0_B_0_, A_0_B_1_, A_1_B_0_, and A_1_B_1_, respectively) were created by subcloning a gBlocks gene fragment (Integrated DNA Technologies) with the point mutation into the wildtype A_0_B_0_ full-length plasmid using KasI-AscI restriction-ligation. Subsequently, the mutation was subcloned from the resulting plasmid containing into the other isoforms’ full-length or ectodomain variants using AgeI-AscI restriction-ligation. Alternatively, by subcloning the regions containing splice inserts using EcoRI-AgeI restriction-ligation into a plasmid already containing the mutation.

### Clustering assay

To introduce plasmids into the cells used for the clustering assays, we employed electroporation as previously reported^29^. K562 cells (catalog nr. ACC 10, Leibniz Institute DSMZ) were cultured in RPMI-1640 medium (Gibco), supplemented with 10% FBS (Gibco) and 1% Penicillin/Streptomycin (Gibco) in a shaking incubator at 37°C and 5% CO_2_. Prior to electroporation, K562 cells were centrifuged for 5 min at 300 g and washed in 1x PBS (Gibco). Cells were again centrifuged for 5 min at 300 g and resuspended in buffer R (Gibco). Per condition, 2 × 10^6^ cells were incubated with a total amount of 15 µg of DNA (Ten3:empty vector ratio was 1:5) for 15 min at room temperature. After the incubation, K562 cells were electroporated with the Neon Transfection System (Thermo Fisher Scientific), using the following parameters: 1450 V, 10 ms pulse length, and 3 pulses^22^. Cells were directly plated onto 5 mL of pre-warmed RPMI-1640 medium with 10% FBS (without Penicillin/Streptomycin) in 6-well plates. Cells were allowed to recover for roughly 20 hours in a shaking incubator at 37°C and 5% CO_2_ before performing the cell clustering assay as described in Chataigner *et al* 2022 ^62^. After recovery, cells were collected and centrifuged for 3 minutes at 200 g. Cells were then resuspended in clustering medium (RPMI-1640 supplemented with 10% FBS) and treated with DNase I (Invitrogen) for 10 minutes at 37°C. Cells were once again centrifuged, resuspended in clustering medium and passed through a 40 μm cell strainer. Cells were counted using the Countess 3 FL (Thermo Fisher Scientific), and a total of 2 x 10^5^ cells per clustering condition were plated in a 12-well plate in 1 mL of clustering medium. Cells were left to cluster for 24 hours on a shaking incubator at 37°C and 5% CO_2_ and clusters were imaged on an EVOS M5000 microscope with a 10x (0.25 NA; EVOS, Thermo Fisher Scientific) or 40x (0.65 NA; EVOS, Thermo Fisher Scientific) objective, using the EVOS GFP light cube (Thermo Fisher Scientific). For analysis, regions of interest (ROIs) larger than 100 pixels were selected after Triangle thresholding of the Gaussian blurred GFP channel images in Fiji^63^. The *Analyze Particles* command was performed to extract the area of each ROI. A cell cluster was defined as an object 3.25X the mean large single cell size (800 pixels)^29^. The clustering index was determined as the summed cluster area divided by summed area of all ROIs within an image (clusters + non-clusters) times 100%. The cluster sizes were averaged per image, and data was acquired from four independent experiments (5 images per experiment, 20 images total per condition). Statistical significance was determined by performing a two-way ANOVA followed by a Šìdák’s multiple comparisons test. All analysed data are represented as mean ± s.e.m.

### AlphaMissense

AlphaMissense^40^ pathogenicity data for human teneurin-3 (hTen3, Q9P273) was acquired from the AlphaFold structure Protein StructureDatabase^64^. Structural data visualisation was performed in PyMOL^65^. AlphaMissense annotated PDB hTen3 data was downloaded from the Hege Lab database, and colour values were visualised in PyMOL using the Hege Lab coloram.py script.

### Protein purification

The mutant variants of the Ten3 isoform ectodomains were purified as reported previously for the wildtypes^29^. Shortly, all Ten3 ECD variants were expressed using Epstein–Barr virus nuclear antigen I-expressing HEK293 cells (HEK-E; U-Protein Express), cultured in FreeStyle293 expression medium with GlutaMAX (FreeStyle; Gibco) supplemented with 0.2% fetal bovine serum (FBS, Gibco) and 0.1% Geneticin (G418 Sulfate; Gibco). Cells were transfected with a total of 125 μg DNA encoding the full ECD using polyethylenimine (PEI, 1:3 DNA:PEI ratio; Polysciences) according to the manufacturer’s protocol and treated with 5.5% Primatone in Freestyle medium 6-24h post-transfection. Medium was collected after six days, and mutant proteins were purified by Ni-NTA affinity chromatography using an elution buffer containing 25 mM HEPES (pH 7.8), 500 mM NaCl, 2 mM CaCl_2_, and 500 mM Imidazole, followed by size-exclusion chromatography (SEC) using a Superose6 Increase 10/300 GL column (Cytiva) into a final buffer composition of 20 mM HEPES (pH 7.8), 150 mM NaCl, and 2 mM CaCl_2_. After each Ni-NTA affinity and SEC purification step, the proteins were concentrated using a 15 mL Amicon ultra centrifugal filter at 100 kDa cut-off.

### Small-angle X-ray scattering

Prior to the small-angle X-ray scattering (SAXS) experiments, protein solutions were shortly centrifuged and final protein concentrations were measured after dilution through absorbance at 280 nm extinction coefficient correction. SAXS measurements were performed at the European Synchrotron Radiation Facility at the BM29 BioSAXS beamline, equipped with the Pilatus3 2M detector. The sample-to-detector distance was fixed at 2.87 m and the X-ray energy was of 12.5 keV, resulting in the scattering vector q range of 0.055 to 4.96 nm^-1^. Three concentrations (targeted at 0.25, 0.5, and 1 mg/mL, see Table 1 for actual concentrations) for each of the Ten3 splice variants and one sample buffer (150 mM NaCl, 20 mM HEPES, 2 mM CaCl_2_, pH 7.8) were loaded into vials of the automatic sample changer, followed by injection of 50 μL of each sample into the 1 mm-diameter quartz capillary. Both vials and the capillary were maintained at 20 °C. Data were collected with 10 frames per sample and a total exposure time of 2 seconds. Buffer frames were collected before and after each sample run. A continuous flow mode was applied to the capillary to minimize the radiation damage. The initial processing of the 2D raw data, including radial averaging, normalization to sample thickness and transmission, and calibration to the absolute intensity in cm^−1^, was performed automatically using the EDNA pipeline^57^, generating 1D I(q) profiles. The frames averaging, buffer subtraction, and further data analysis were then performed by BioXTAS RAW software^61^. The frames with aberrant scattering (e.g. showing radiation damage at low q) were manually excluded from the averaging. The averaged and buffer-subtracted I(q) data were then normalized by the protein concentration for each sample. The 1 mg/mL data, with the best signal-to-noise ratio, were chosen for further analysis, with exception for A_0_B_1_ wildtype of 0.5 mg/mL. The A_0_B_1_^MCOP^ and A_1_B_0_^MCOP^ curves of 1 mg/mL were merged with the corresponding 0.25 mg/mL ones at the lowest q (up to 0.113 nm^-1^ and 0.13 nm^-1^, respectively), due to slight inter-particle effect visible at the lowest q. The scattering data were analysed by direct modelling with Guinier approximation, to determine the radius of gyration (R_g_) and the forward scattering intensity, I(q→0), through linear fitting of ln I(q) versus *q*^2^. The regions of the Guinier fits were chosen for sufficiently small angles, i.e. q·Rg ≤ 1.3. The real-space pair distance distribution function, P(r), with maximum dimension (D_max_) was calculated by running GNOM program with Indirect Fourier Transform of the I(q) curves to 3 nm^-1^ truncated^59,60^. The molecular weight (M_w_) estimates were calculated through I(0), Porod volume (V_p_), and the Bayesian inference^66^. The Porod-Debye transform (I(q)q^4^ vs. q^4^) was used to identify the Porod regions for calculating V_p_. Dimensionless Kratky transform (qR_g_)^2^I(q) / I(0) vs. qR_g_) was used for structural insights.

### Molecular dynamics simulations

The starting structures for the molecular dynamics simulations were based on the protein structure of the Ten3 compact dimer (PDB: 8R50^29^). Glycan chains were removed from the structure. Unsolved gaps in the protein structure were modelled using Alphafold 3^67^. The starting positions for the EGF domains were modelled based on SAXS experiments, as described in Gogou *et al.* 2024^29^. The structure was capped at both the N- and C-termini with a methyl and acetyl group, respectively, using the capping functionality of PyMOL^65^.The mutagenesis wizard of PyMOL was used to introduce the R2579W mutation and generate the mutant Ten3 dimer. PDB2PQR^68^ was used to generate the protonation state corresponding to a pH of 7.8. In addition, six structural calcium ions were added, which are likely present in teneurin-3 due to sequence conservation (PDB: 7PLP^34^), which remained in place throughout all simulations without additional restraints. Simulation of the wildtype and mutant version of the Ten3 dimer were run in triplicate, yielding six separate simulations.

Parameterization of the Ten3 structures was carried out using the *tleap* tool of AmberTools23, version 23.6^69^. We used the ff14SB force field^70^ combined with the TIP3P water model^71^ and Li and Merz Na^+^, Cl^-^ and Ca^2+^ ions^72,73^. Known disulfide bridges were introduced at previously identified sites (UniProt: Q9WTS6). Finally, ParmEd 4.1.0^74^ was used to convert the file format from Amber to GROMACS.

Subsequent minimization, equilibration, production, post-processing and analysis was carried out using the GROMACS software suite, version 2023.3^75–81^. The energy of each system was minimized using the steepest descent algorithm until a maximum force of 1000 kJ mol^-1^ · nm^-1^ was reached. The equations of motion were numerically integrated using the Verlet leapfrog algorithm with a timestep of 2 fs. Coordinates for each atom were written every 10 ps. A cutoff of 1 nm was used for short-range electrostatic and van der Waals interactions, whereas the long-range electrostatic interactions were calculated by particle-mesh Ewald summation with fourth order cubic interpolation and a grid spacing of 0.16 nm^82^. After minimization, the systems were equilibrated for 100 ps in the *NVT* ensemble using the modified Berendsen thermostat to reach the equilibrium temperature of 310 K. Next, the system was equilibrated in the *NpT* ensemble, for 100 ps, using the Parrinello-Rahman barostat to reach and equilibrium pressure of 1.0 bar^83^. The minimized and equilibrated systems were then produced for 1 μs each, in the *NpT* ensemble, during which all hydrogen-containing bonds were constrained using the LINCS algorithm^84^.

The obtained trajectories were corrected for periodic boundary conditions (pbc) using the *whole* and *nojump* options of *gmx trjconv -pbc*. For visualization purposes, we trimmed the full precision trajectories by taking a stride of either 100 frames or 1000 frames and rotationally and translationally fitted each frame to the ABD domains using *skip* and *fit* flags of *gmx trjconv*.

The binding affinity between the two subunits was obtained through the Molecular Mechanics energy with General Born and Surface Area continuum solvation method (MM/GBSA), as described in^85,86^. We used the last 500 ns of each trajectory, taking 500 frames, to perform the calculations. The gmxMMPBSA tool was used to more easily apply the Amber MMPBSA.py method to the trajectories, which were generated in GROMACS^87^.

The Visual Molecular Dynamics (VMD) S*alt Bridges* plugin was used to obtain the salt bridges in the simulations. We used the trajectories with a stride of 1000 and a cutoff distance of 4 Å. All acidic (D, E) and basic (R, H, K) residues are considered. The normal modes were obtained through the *Normal Mode Wizard* embedded in VMD^88^. We performed an ANM calculation, using the Cα atoms of the dimer without the EGF domains, a cutoff distance of 15 Å and a unit force constant.

The mutation site contacts, r.m.s.d., r.m.s.f., radius of gyration and hydrogen bonds were obtained by employing GROMACS commands, namely (gmx) *select*, *rms*, *rmsf*, *gyrate* and *hbond*. For the contacts we chose a cutoff distance of 4 Å and select all atoms. The r.m.s.d. and r.m.s.f. were calculated using the backbone atoms of the dimers. The radius of gyration was calculated using all atoms in the protein and the hydrogen- and ionic bonds were filtered for interchain hydrogen bonds.

### Thermal stability assay

Thermal shift assay (TSA) assays were performed as previously reported^29^ using purified human Ten3 isoforms and their MCOP mutants at a concentration of 0.5 mg/mL in SEC buffer (20 mM HEPES, 150-mM NaCl, 2 mM CaCl_2_, pH 7.8). Melting curves were measured of five technical repeats for each sample. The melting temperatures (T_m_) were determined as the maximum of the first derivative of the fluorescence signal.

### Automated SEC(-MALS)

For analytical size exclusion chromatography (aSEC), Ten3 isoforms, along with a bovine serum albumin (BSA) reference, were diluted to a final concentration of 0.5 mg/mL in SEC buffer (20 mM HEPES, 150 mM NaCl, 2 mM CaCl_2_, pH 7.8) and automatically loaded onto a Superose6 Increase 10/300 GL column (Cytiva) integrated with a high-performance liquid chromatography (HPLC) unit (1260 Infinity II, Agilent) coupled to an online UV detector (1260 Infinity II VWD, Agilent) and a refractometer (Optilab T-rEX, Wyatt Technology) in series. For the undigested proteins, the experiment was repeated twice in opposite order, to exclude the possibility that differences in run time arose due to the waiting times in the sample holder before running. For the Ten3 A_1_B_1_ wildtype variant, two technical were consecutively run in one experiment to determine the degree of reproducibility for identical samples in direct succession. To fully strip all glycan chains of the Ten3 surface, digestion with PNGase F (New England Biolabs) was performed. Buffer exchange by 7K MWCO Zeba spin desalting columns (Thermo Fisher Scientific) was first performed on the PNGase to remove the glycerol from the storage buffer. 50,000 U of PNGase in SEC buffer buffer was then added to a final 50 μL of 0.5 mg/mL protein to incubate overnight at 17°C. The next day, the temperature was increased to 30°C for another 2 hours of digestion and the sample was frozen in liquid nitrogen for storage in −80°C. Before running on aSEC, the sample was thawed on ice and centrifuged for 10 minutes at 11,000 g and 4°C to remove aggregates.

Purified proteins were diluted to a final concentration 20 µg/mL, boiled for 10 min at 98 °C in the presence or absence of β-mercaptoethanol and loaded onto a TGX mini protean precast gel (Biorad) for 70 min at 200V.

### Cryo-EM data acquisition and single-particle analysis

Cryo-EM imaging of Ten3 A_1_B_1_ R2579W mutant ECD was performed on purified protein diluted in SEC-buffer supplemented with 2 mM CaCl_2_ to a final protein concentration of 0.5 mg/mL. 3.0 μL of diluted sample was deposited onto Quantifoil R1.2/1.3 Cu 300 mesh grids that were glow-discharged. Excess sample volume was blotted away for 2.5-4.0 seconds with blot force of −3.0 at 22°C and 100% humidity before plunging into liquid ethane using a Vitrobot Mark IV (Thermo Scientific). Movies were acquired using a K3 detector (Gatan) in counting super-resolution mode with a 20 keV (Gatan) slit width at 105,000X nominal magnification, a corresponding super-resolution pixel size of 0.418 Å, and a total dose of 50 e^-^/Å^2^ per movie. A total of 13,326 movies were acquired at a defocus range of −0.8 to −2.0 μm. Beam-induced motion and drift correction was performed using MotionCorr2^89^ with a pixel binning factor of 2 (0.836 Å physical pixel size) in RELION4.0^90^. CTF estimation using RELION’s CTFFIND^91^, particle picking, and first rounds of 2D- and 3D classification were performed in RELION5.0 on CryoCloud. Particles were initially picked using RELION’s 3D reference-based picking, using a 20 Å low-pass-filtered compact dimer reference from the Ten3 A_1_B_1_ wildtype^29^. In parallel, reference-free particle picking was performed using CryoCloud picker, upon which the picking jobs were joined and duplicated particles were removed. 2D classification was used to discard non-particle objects like foil hole edges and ice crystals from the extracted particle sets. For all datasets, 3D map reconstructions were performed according to the regular workflows of Relion4.0, and as reported in the corresponding Supplementary Fig. 4. Compact dimer particles were first isolated using a masked wildtype compact dimer as a reference for 3D classification. The resulting particle set comprising both open and closed conformations were converted into a particle stack at 2-fold downsampling for 3D variability analysis in CryoSPARC^92^. Multiple rounds of 3D Classification with and without alignment were performed in RELION4.0 to further isolate the closed conformations, after which iterative particle polishing and CTF refinement steps were performed to attain the highest-resolution reconstruction with C2 imposed symmetry. For reconstruction of the non-compact subunit, a non-compact subunit reference was used from the wildtype pipeline^29^. After first performing single-class 3D classification with a mask around the full non-compact particle (TTR domain through Tox-GHH domain), iterative 3D classification without alignment was performed with a mask around the YD, ABD, and Tox-GHH domains to start selecting for particles good density for the ABD domain. To reduce calculation times due to the many remaining particles, a random subset of 30% was selected before continuing with particles subtraction around the same YD-ABD-Tox-GHH mask. Before particle subtraction, the particles were polished, and particles were CTF refined after subtraction. Subsequently, focused 3D classification without alignment using a mask around the ABD and Tox-GHH domains, and 3D refinements were iteratively used to select particles to achieve an optimal reconstruction of the ABD domain’s solvent-exposed β-sheet. Protein sequence alignments of the calcium binding sites across teneurin mouse and human family members were generated using Clustal Omega^93^.

### Model building and refinement

Wildtype compact dimeric Ten3-A_1_B_1_ (PDB:8R50) was combined with an Alphafold 3^67^-predicted model for the region containing R2579W (residues 2570-2580) that was unstructured in the wildtype. The resulting model was refined in the MCOP compact dimer in closed conformation by iterative cycles of phenix.real_space_refine of the Phenix package^94^ and manual real-space refinement in Coot^95^. During these steps, missing residues that were present in the map, such as I747, were built into the model and refined during these iterations. The stereochemistry of the models was checked with MolProbity^96^. To calculate the r.m.s.d. value between previously published wildtype (PDB:8R50) and the novel MCOP (PDB:9IGS) compact dimers, PyMOL’s^65^ *align* function was used. To quantify experimental differences between the 3D variability analysis of wildtype and mutant compact dimer movements, the single subunits of the compact dimer were rigid body fitted into the volumes of the fully closed and fully open states. The resulting compact dimer structures were then again compared by calculating the r.m.s.d. using PyMOL’s *align* function. The figures containing structural maps and models were generated using Chimera^97^ and PyMOL.

## Data availability

The SAXS data generated in this study have been deposited in the Small Angle Scattering Biological Data Bank (SASBDB) under the accession codes SASDW78, SASDWY7, SASDWZ7, SASDWQ7, SASDWR7, SASDW88, SASDW98, SASDWS7, SASDW28, SASDWT7, SASDWA8, SASDWB8, SASDW38, SASDW48, SASDWU7, SASDWV7, SASDWC8, SASDW58, SASDW68, SASDWW7, SASDWX7.

The structure model data generated in this study has been deposited in the Protein Data Bank (PDB) under the accession codes 9IGS (teneurin-3 A_1_B_1_ MCOP compact dimer). The cryo-EM density map data generated in this study have been deposited in the Electron Microscopy Data Bank (EMDB) under the accession codes EMD-52854 (teneurin-3 A_1_B_1_ MCOP compact dimer) and EMD- 52855 (teneurin-3 A_1_B_1_ MCOP non-compact subunit).

## Code availability

Scripts and code used for K562 clustering analysis in this study are used as previously described in Gogou *et al.* 2024^29^ and are available through Zenodo [https://doi.org/10.5281/zenodo.10843181].

## Inclusion and diversity statement

In line with our commitment to promoting a diverse and inclusive scientific community, we affirm our dedication to upholding the principles of equity and diversity in the research presented in this paper. We believe that diversity in the scientific community is essential to drive innovation, foster creativity, and ensure comprehensive and unbiased exploration of the life sciences.

## Acknowledgements

We would like to acknowledge Stephany Laclé for her contribution to preparing the molecular dynamics simulation. We would like to acknowledge Cecilia de Agrela Pinto for her assistance and guidance in performing and analyzing the analytical size-exclusion chromatography experiments. We would like to thank Mike Filius for designing the PNGase digestion experiment and for kindly providing the enzyme.

We acknowledge the European Synchrotron Radiation Facility (ESRF) for provision of synchrotron radiation facilities under proposal number MX-2707, and we would like to thank Mark Tully for assistance and support in using beamline BM29 for data collection (10.15151/ESRF-ES-1984114240). Cryo-EM data were collected at The Netherlands Centre for Electron Nanoscopy (NeCEN) with assistance from Ludovic Renault. RELION5.0 was used on CryoCloud. Molecular dynamics simulations were in part executed on DelftBlue Supercomputer^98^. DHM is supported by an NWO computing grant (2021.058) and NWO Veni grant (722.016.004).

## Author contributions statement

Conceptualization: CG, DHM

Methodology: CG, SL, MN, CPF

Data acquisition and analysis: CG, SL, MN, CPF

Supervision: DHM, NS

Writing – original draft: CG, DHM

Writing – review & editing: All

## Competing interest declaration

The authors declare no competing interest.

## Notes

### Competing Interest Statement

The authors have declared no competing interest.

