## Supplementary Figures for "A teneurin-3 microphthalmia mutation disrupts *trans* adhesion for specific alternative splicing isoforms"

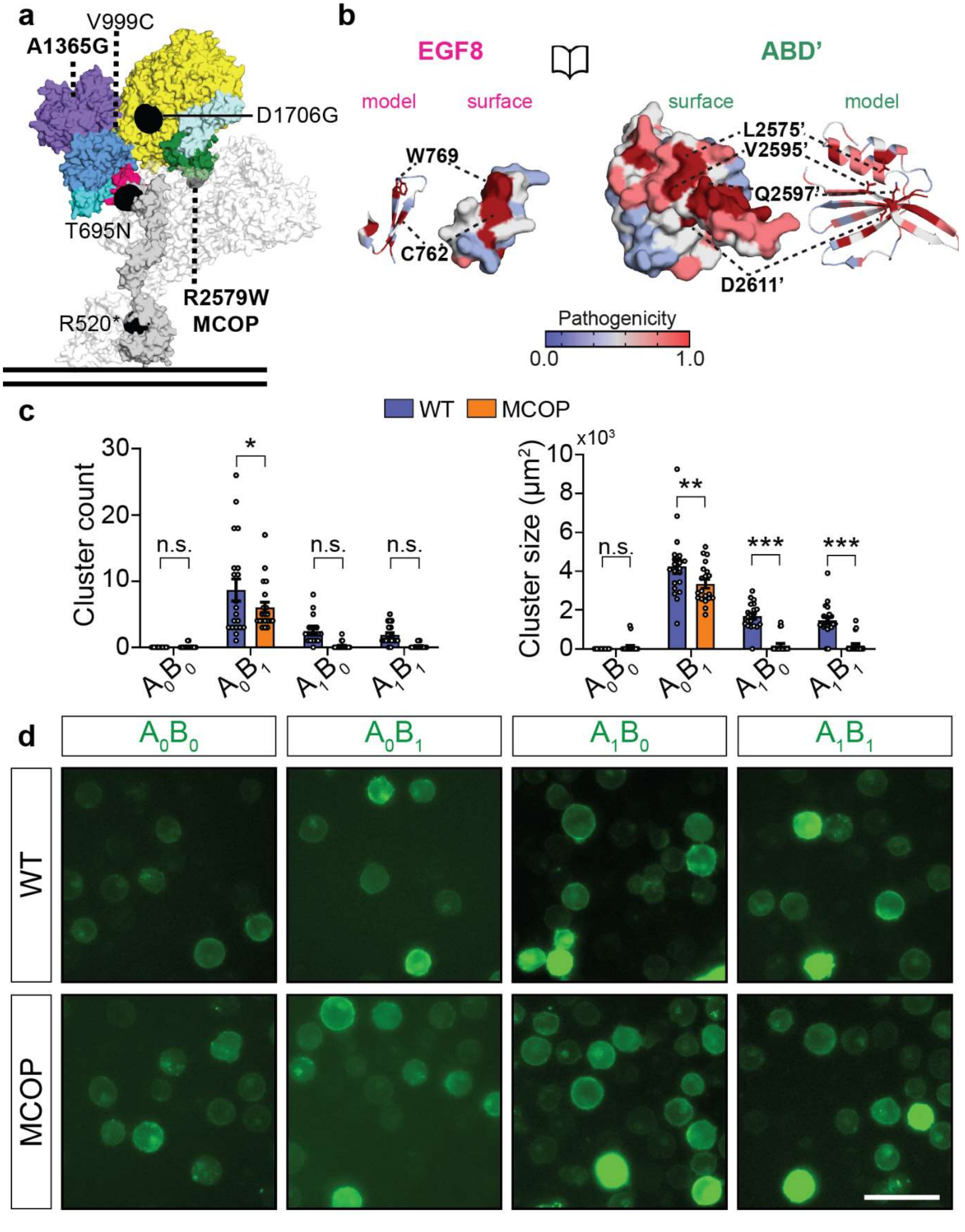


Supplementary Figure 1: R2569W is located at the EGF8-ABD contact and does not interfere with expression or trafficking. **a)** Surface representation of mouse Ten3 ECD in $A_1$ compact dimer configuration. One covalently linked monomer is coloured corresponding to Fig. 1b, and the other is shown as transparent white surface. Black annotated spheres represent the missense mutations form Fig. 1a. Black dashed lines indicate mutations that are buried in the structure. R2579W is buried between the coloured and transparent subunits. Vertical black lines represent the membrane. The unique compound heterozygous missense mutations are indicated in bold in all panels. **b)** Open book surface representation of AlphaMissense-predicted pathogenicity at the EGF8-ABD interface. Corresponding models are shown beside the surface representations. Hydrophobic, as well as hydrogen- and ionic bond-forming interface residues are labelled and indicated with black dashed lines. Residue numbering corresponds to the $A_1B_1$ isoform. The pathogenicity score colour coding legend is shown in the bottom right. **c)** Additional quantification of clustering data in panel Fig. 1d. Bar plots represent the number of clusters found and the average sizes of the clusters per image ($n$=5 images per $N$=4 independent experiments, 20 images total). Data are presented as mean ± s.e.m.; ns: not significant, *: p > 0.05, **: p > 0.01, ***: p > 0.001. Two-Way ANOVA, Šìdák's multiple comparisons test. Specific p-values from left to right are for cluster count >0.999, 0.025, 0.067, and 0.248, and for cluster size 0.988, 0.003, <0.0001, <0.0001. **d)** Representative fluorescence microscopy images of non-clustering K562 cells at 40x magnification expressing the wildtype (WT) Ten3 isoforms and microphthalmia mutants (MCOP). Scale bar is 50 µm.

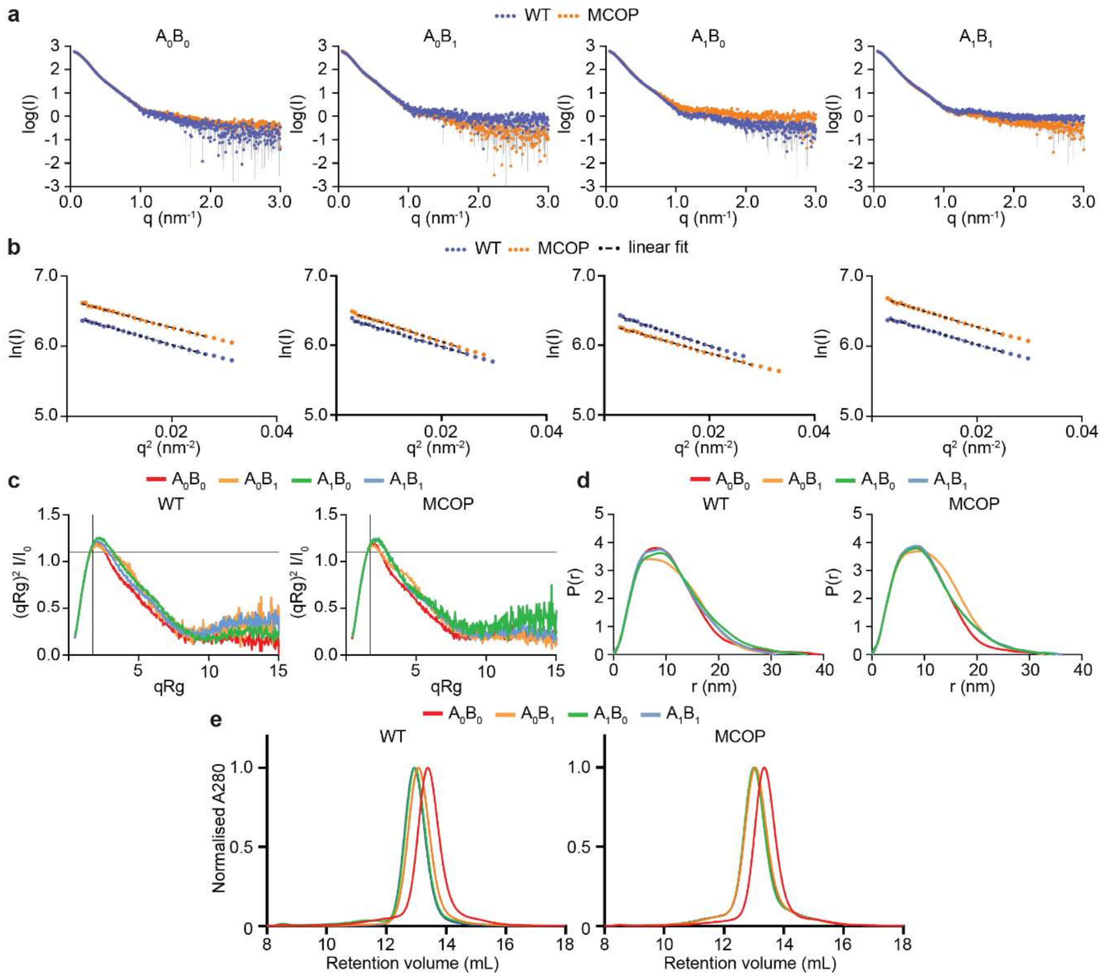


Supplementary Figure 2: Size-exclusion chromatography (SEC) and small-angle X-ray scattering (SAXS) analysis of all Ten3 isoforms and their respective MCOP mutants. **a)** SAXS log I(q) vs. q profiles for all purified Ten3 isoforms and their respective MCOP mutants. **b)** Guinier plots with corresponding linear fits for all purified Ten3 variants in panel a. the range of Guinier fits was chosen for q.Rg ≤ 1.3. See Table 1 for supporting the quality of the fits. Values for $A_0B_0$ and $A_1B_1$ MCOP mutants were shifter by arbitrary value 0.25 to visualize distinguish individual plots. **c)** Comparison of all wildtype (WT, left) and all mutant (MCOP, right) dimensionless Kratky plots with maxima of ~1.1 at $qR_g$ of √3 (marked with the crossings of the lines), indicating a geometry of a globular particle[1]. **d)** Comparison of all WT (left) and all MCOP (right) pair distance distribution functions. As seen in Table 1, the $R_g$ and $D_{max}$ of $A_1B_1$ increase upon introduction of the MCOP mutation, thereby becoming more similar to the values of the $A_1B_0$ wildtype and mutant. Also, in the Kratky plots and P(r) presented here, the $A_1$ isoforms become much more similar in overall structure in the presence of the mutation. **e)** Comparison of all WT (left) and all MCOP (right) analytical SEC runs.

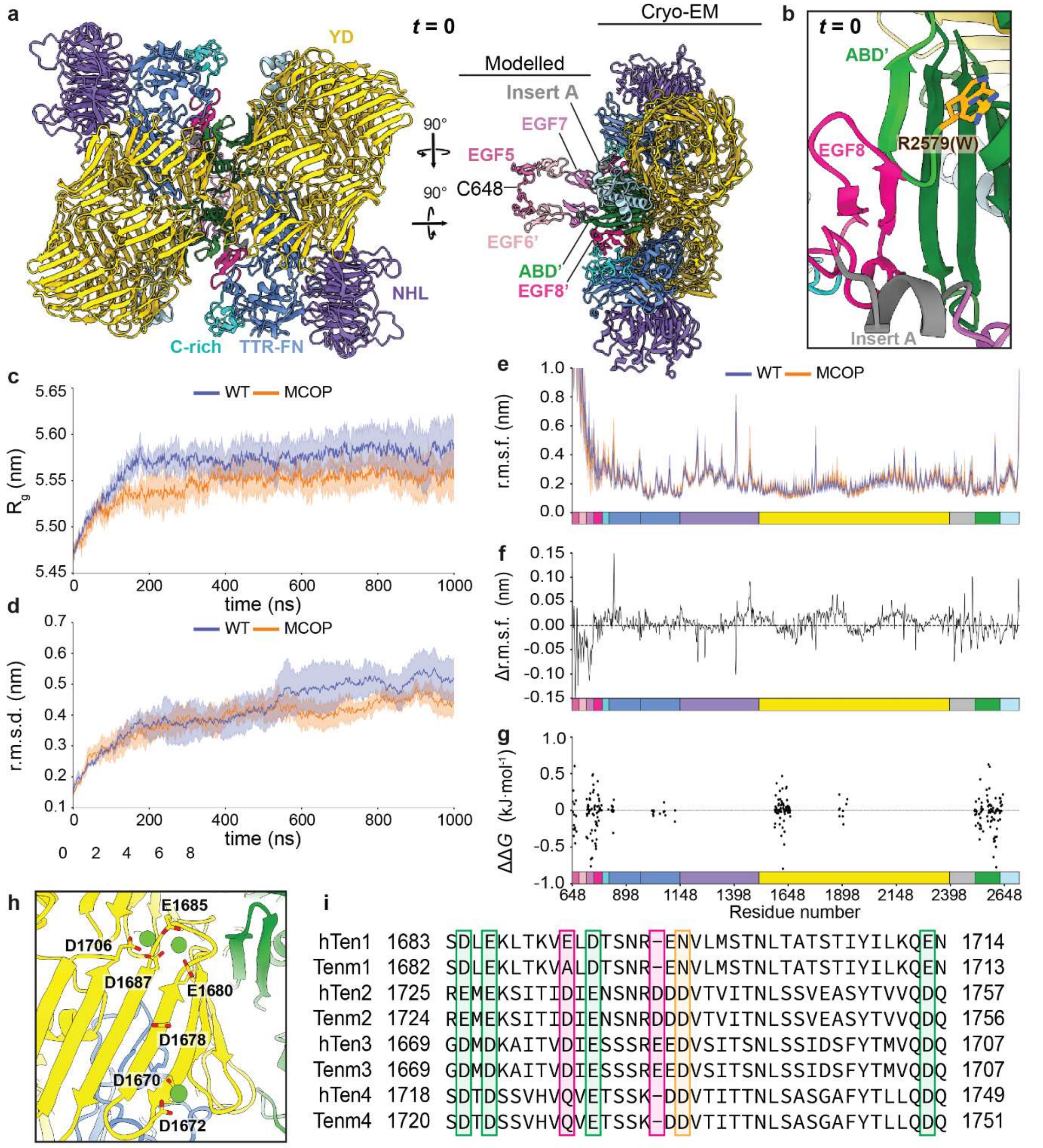


Supplementary Figure 3: Molecular dynamics simulations starting structure and additional data. **a)** Complete view of the MD simulations' starting model. The WT compact dimer structure (PDB:8R50) is supplemented with EGF5-EGF7, derived from the SAXS-modelled extracellular domain[2]. The most N-terminal cystine (C648) of the simulated system is indicated, which also covalently links the two monomers. **b)** Inset showing the single-residue difference between the WT (R2579) and MCOP (R2579W) starting models (at $t = 0$). The wildtype arginine and mutant tryptophan are both displayed. The light green region indicates the loop and flanking β-strand that are originally unstructured in the cryo-EM compact dimer. Domain colour codes of panels c-e correspond to the description in Fig. 1b, and apostrophes (') are used to distinguish between residues located on one subunit in the compact dimer vs. localization on the other subunit. **c)** Time traces of the radius of gyration ($R_g$) for WT and MCOP molecules. **d)** Time traces of the r.m.s.d. relative to the starting structure for WT and MCOP molecules. **e)** Per-residue r.m.s.f. relative to the average position in a trajectory for WT and MCOP molecules. Data in panels c-e represent mean ± s.e.m. of three simulations per variant. **f)** Difference in per-residue r.m.s.f. plots shown in panel e as r.m.s.f. of MCOP minus r.m.s.f. of WT. **g)** Difference in per-residue binding free energy ($\Delta G$) between WT and MCOP mutant, calculated as MCOP minus WT. **h)** Region on the YD shell that was consistently bound by calcium ions throughout all WT and MCOP simulations. **i)** Conservation analysis of the calcium-binding residues identified in panel h through sequence alignment of human teneurin family members (hTen) and mouse teneurin family members (Tenm). Green boxes indicate preservation of negatively charged residues; a red box indicates missing residues in some of the orthologues; orange boxes indicate substitution of a negative charge by an uncharged residue.

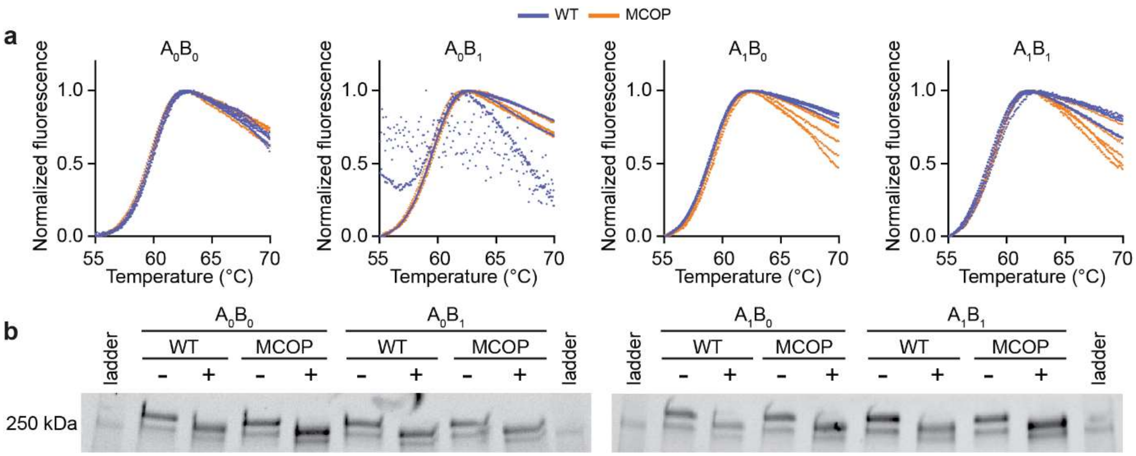


Supplementary Figure 4: Thermal stability assay in the absence of calcium and PAGE validation of PNGase digestion. **a)** Melting curves of all Ten3 isoforms and their respective MCOP mutants in the presence of 5 mM EDTA. All five technical repeats per variant are plotted. **b)** PAGE gels of all Ten3 variants with (+) and without (-) PNGase digestion to strip all glycan chains from the ectodomains. All PNGase-treated samples end up lower on gel, indicating successful stripping of the high-molecular weight glycans. The 250-kDa band from the ladder is visible in all ladder lanes but only annotated on the outermost left lane.

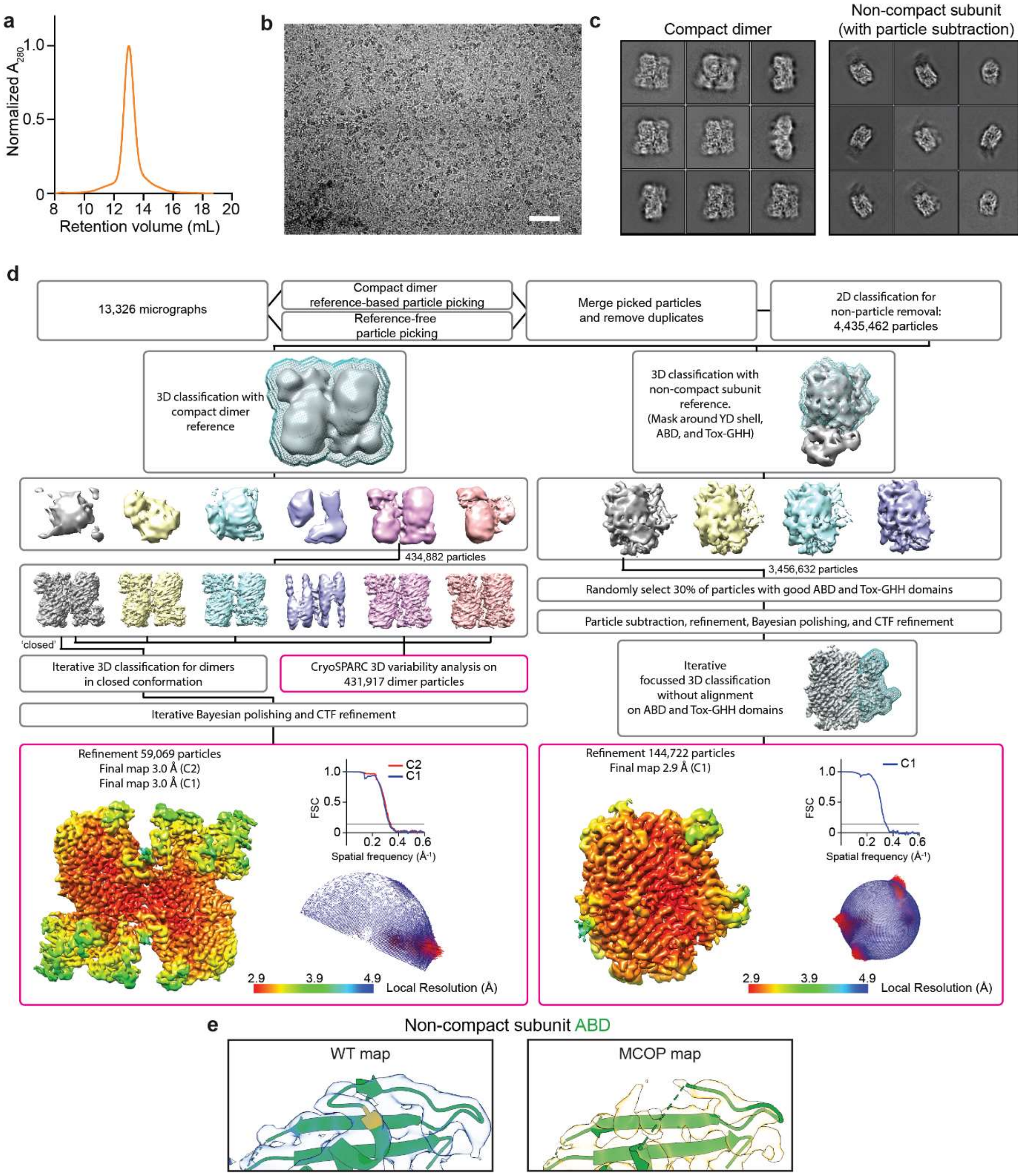


Supplementary Figure 5: Cryo-EM data collection and single-particle analysis (SPA) of Ten3-$A_1B_1$ R2579W (MCOP). **a)** SEC trace of the Ten3-$A_1B_1$ MCOP ectodomain after nickel affinity purification**.** The same data is plot as in Supplementary Fig. 2e. **b)** Representative cryo-EM micrograph of vitrified Ten3-$A_1B_1$ MCOP ectodomain. Scale bar is 50 nm. **c)** SPA 2D classes of compact dimeric Ten3-$A_1B_1$ MCOP particles in 'closed' conformation (left) and of non-compact subunit after particle subtraction outside YD through Tox-GHH domains (right). **d)** SPA workflow for the reconstruction of cryo-EM electron densities of Ten3-$A_1B_1$ MCOP. Left: reconstruction workflow of the compact dimeric map. Right: focussed classification and refinement after subtraction of the non-compact subunit. Pink outlined boxes denote endpoints of the SPA workflow. Corresponding local resolution values, Fourier shell correlation (FSC) graphs, and angular distribution of particles in final refinements are shown for each map. FSC cut-off is 0.143. Source data for panels A and D are provided as a Source Data file. **e)** Comparison of the WT (EMD-18890) and MCOP (EMD-52855) ABD domain density in the cryo-EM-reconstructed non-compact subunits. The region that could not be resolved in the MCOP mutant density was manually removed from the fitted WT non-compact subunit (PDB:8R51) and corresponds to the region that is not resolved in the WT EGF8-ABD compact dimer interface. Domain colouring corresponds to the description in Fig. 1b. The mutation site at position 2579 that was resolved in the WT but not in the mutant is coloured orange.

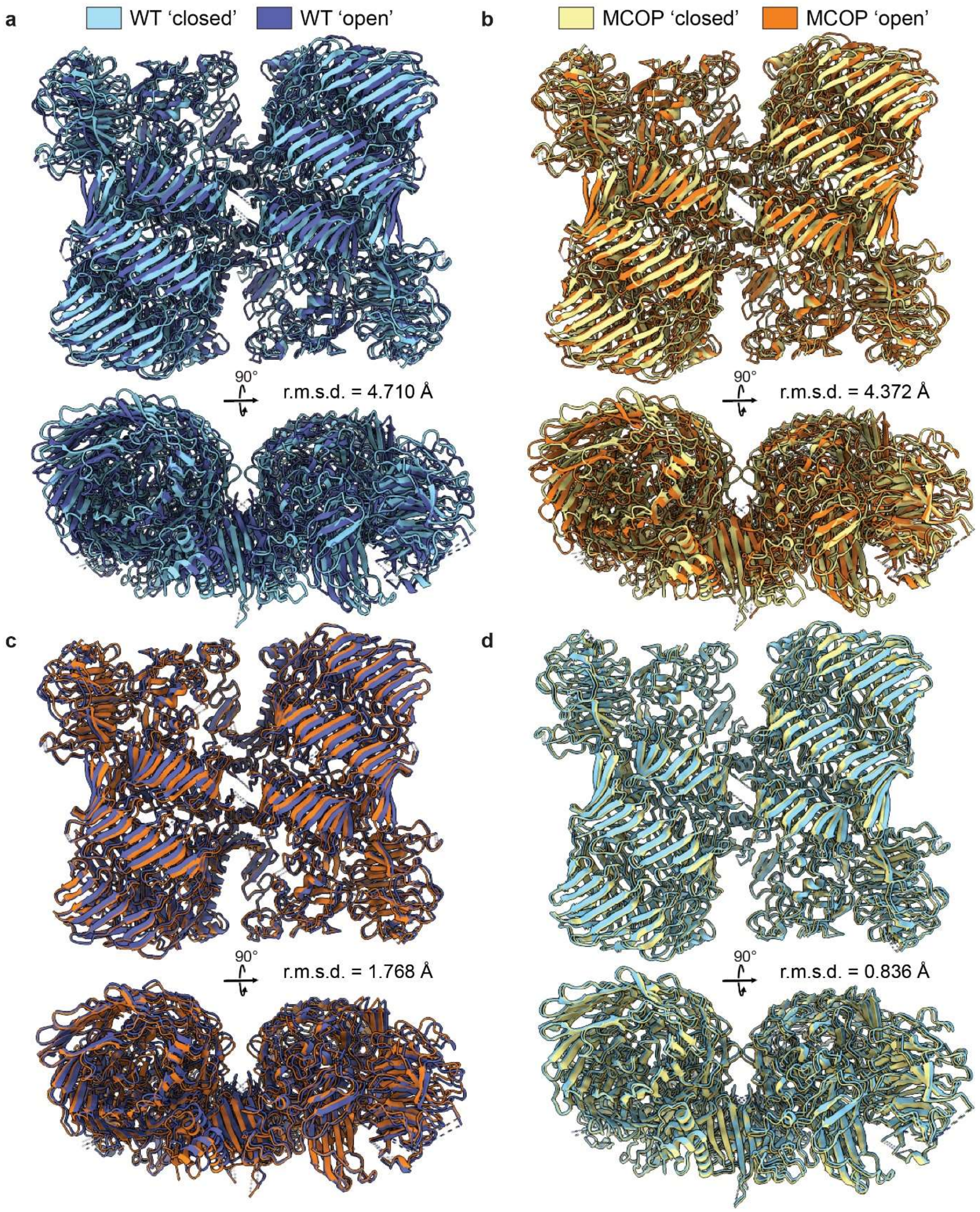


Supplementary Figure 6: Comparison of the open and closed states of the wildtype and mutant butterfly movements. **a)** Overlay of the wildtype (WT) open and closed conformation structures. **b)** Overlay of the mutant (MCOP) open and closed conformation structures. **c)** Overlay of the WT and MCOP open conformation structures. **d)** Overlay of the WT and MCOP closed conformation structures. For the pairs of overlayed structures in each panel, the r.m.s.d. value is shown. Front and top views of each overlay are displayed. The closed conformations are recognizable in the top views where the hydrophobic loops in the YD shells of the two subunits connect. All glycans were hidden in these structures for visualization purposes.

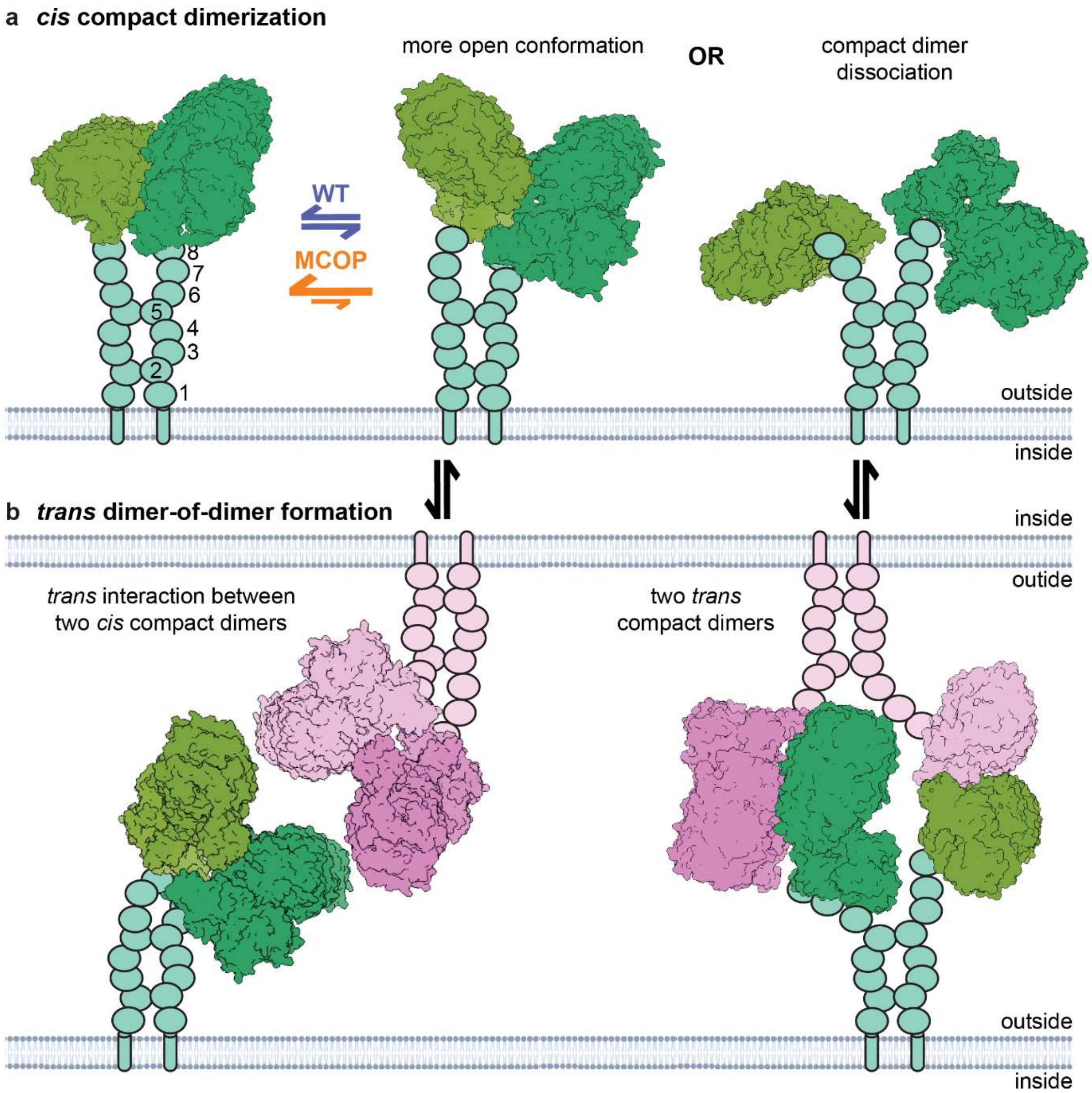


Supplementary Figure 7: Extended model for MCOP-induced disruption of *trans* adhesion via stabilization of the *cis* compact dimer. Schematic visualizations of a Ten3 $A_1$ covalent *cis* dimer (green) attached to the extracellular side (outside) of membrane. Left and middle: A WT compact dimer in *cis* has more conformations freedom to allow changes the relative orientation of its subunits, whereas the mutation restrict the conformational landscape of the duplex. Right: Representation of the two subunits completely dissociating the compact dimer interface, leaving them only attached by the disulfide links in EGFs repeats 2 and 5. b) Formation of *trans* complexes between covalent dimers on two opposite membranes. Left: The higher conformational freedom of the wildtype allows the compact dimers in cis to adopt a conformation that allows an interaction with another compact dimer (pink) in *trans*. Right: Dissociation of the cis compact dimer in panel a is an intermediate step before each subunit can form a new compact dimer interface with the subunit from another covalent dimer in *trans*. We thus hypothesize that stabilization by the MCOP mutation the conformational changes in panel a required to form the *trans* dimer-of-dimers shown in panel b. All Ten3 molecules are attached to the extracellular side of the membrane via and two sets of a transmembrane helix (tube), an Ig domain (rectangle), and eight EGF repeats (ovals) in the same colour as the attached subunits. Intracellular (inside) domains are omitted.
